# Use of immunoglobulin-binding bacterial proteins in immunodetection

**DOI:** 10.1101/2020.10.08.331462

**Authors:** Angel Justiz-Vaillant, Belkis Ferrer-Cosme, Albert Vera

**Affiliations:** Department of Para-clinical Sciences. Faculty of medical Sciences. University of the West Indies. St. Augustine. Trinidad and Tobago. West Indies.; Immunology Laboratory. Clinical Laboratory. “Saturnino Lora Torres” Provincial Teaching Clinical Surgical Hospital. Province of Santiago de Cuba. Cuba.; Higher Institute of Medical Sciences of Santiago de Cuba. Province of Santiago. Cuba; Immunology Section. Department of Pre-Clinical Sciences. University of Medical Sciences of Cienfuegos. Province of Cienfuegos. Cuba.

**Keywords:** Zoology, bacterial proteins, enzyme-linked immunosorbent assay (ELISA)

## Abstract

One of the aim of this study was to make universal chimeric conjugates to react with both avian and mammalian immunoglobulins in enzyme-linked immunosorbent assays (ELISAs). The periodate method was used in the conjugation process of cross-linking horseradish peroxidase to immunoglobulin-binding proteins (IBP) including staphylococcal protein A (SpA), streptococcal protein G (SpG) and peptostreptococcal protein L (SpL). By mixing up these three conjugates another four hybrid protein conjugates were created including protein LA (SpLA), protein LG (SpLG), protein AG (SpAG) and protein LAG (PLAG). Thirty-five ELISAs were standardized by a probabilistic combination of these immunoreagents. By using a panel of mainly mammalian immunoglobulins their reproducibility was checked by the determination of coefficient of variations (CV) for each one of the IgG-IBP binding. The source of immunoglobulins was their purification by affinity chromatography using a commercially available kit (PURE-1A). The other aim was to immunize chicken with the peptide fragment 254-274 of gp120 to produce anti-HIV peptide hyper-immune egg. Cats and rats were fed these eggs for a determined period until they produced the anti-HIV peptide antibody, which was tested by an indirect SpLA-ELISA and dot blot analysis that corroborated the production of anti-HIV antibodies by the mammalian species including positive humans samples for HIV. We conclude that the single and hybrid immunoglobulin-binding protein were effective in their binding capacity to immunoglobulins from a variety of mammalian species. The potential use of this proteins is in the arena of immunodiagnosis and immunoglobulin detection. Dot blot analysis proves effective in the detection of HIV anti-gp120 antibodies in several animal species. These antibodies can be used as reagents in the development of immunodiagnostic tests or experimental vaccines.

## Introduction

Several bacterial immunoglobulin (Ig)-receptors have been identified. They have proved to be powerful tools for binding, detection, and purification of immunoglobulins [1]. The better studied bacterial Ig receptors include the protein A (SpA) of *Staphylococcus aureus* [2]; protein G (SpG) of *Streptococci* [3,4]; and protein L (SpL) originally isolated from the cell wall of the anaerobic bacterium *Peptostreptococcus magnus* [5].

These bacterial proteins displayed on the cell wall of microorganisms play an important role in bacterial escape mechanisms from the immune system. They cause activation of the complement system by the classical pathway, polyclonal activation of B-lymphocytes, inhibition of phagocytosis and other effects [6–8]. In addition, they have the biological property of binding to a wide range of mammalian and non-mammalian immunoglobulins.

This binding does not interfere with the antigen binding sites on the immunoglobulin receptors. These receptors have been called immunoglobulin-binding protein, IBP [9–14]. Protein A and G have been used as immunological tools in serological tests used in the immunodiagnosis of infectious diseases, such as Borrelia burgdorferi in zoo animals [14]. Ongoing studies suggest that the bacterial Ig receptors are also potential tools in biomedical research, therapy of human diseases, biotechnology, and industry.

Several bacterial immunoglobulins (Ig)-receptors have been identified in recent years. Indeed, they have proved to be robust tools for binding, detection, and purification of immunoglobulins. Among the better-studied bacterial Ig-receptors include the protein A (SpA) of Staphylococcus aureus, protein G (SpG) of Streptococci, and protein L (SpL) originally isolated from the cell wall of the anaerobic bacterium Peptostreptococcus magnus. These bacterial antigens displayed on the cell wall of microorganisms play an essential role in bacterial escape mechanisms from the immune system. They cause the complement system’s activation by the classical pathway, polyclonal activation of B-lymphocytes, phagocytosis inhibition, and other effects [1–5].

Besides, they have the biological property of binding to a wide range of mammalian and non-mammalian immunoglobulins. This binding does not interfere with the antigen-binding sites on the immunoglobulin receptors. They have been called immunoglobulin-binding protein, IBP. SpA and SpG have been used as immunological reagents in serological tests used in the immunodiagnosis of infectious microorganisms, such as Borrelia burgdorferi, in zoo animals. Ongoing studies suggest that the immunoglobulin-binding proteins (IBP) are also practical tools in biomedical research and biotechnology [6–12].

SpA has molecular weight (MW) approximately 42 kDa [1]. It has the capacity to bind to the Fc fragment of IgG produced by many animal species, including humans, dogs, rabbits, hamsters, monkeys, and others [6,13,14]. The native SpA consists of five domains. Of these, four-show high structural homology, containing approximately 58 amino acids and have the capacity of binding to Fc regions of IgG [1].

SpG is a type III bacterial Fc receptor. It is a small globular protein produced by several Streptococcal species and comprises 2 or 3 nearly identical domains, each containing 55 amino acids (aa). SpG binds to the Fc regions of IgG from many mammalian species [15–17].

The molecule contains five homologous “B” repeats of 72-76 aa, and responsible for the interaction with Ig-L chains [18]. SpL is composed of an alpha-helix packed against a 4-stranded beta-sheet [19]. The SpL binds firmly to human kappa light chain subclasses I, III, and IV from the five human Ig classes. Also, SpL binds to other mammalian Ig molecules [5].

In this paper several Immunoglobulins from diverse animal species were purified and used for the standardization of 35 enzyme-linked immunosorbent assays based on immunoglobulin-binding proteins. Their affinity, intra- and inter-assay coefficient of variation were assessed as a measure of the reproducibility of assays, and surprisingly they were all reproducible and sensitive enough for the evaluation of the different affinity of IBP to these antibodies.

## Materials and Methods

A commercial protein-A affinity chromatography called PURE-1A (Sigma-Aldrich.) was used to purify mammalian immunoglobulins from the serum of different mammalian species including horse, mule, dog, coyote, and others. [20]. The instructions of the manufacturer were followed in performing this procedure. Briefly Anti-SpA containing serum is first loaded onto the Protein A Cartridge where the IgG is immobilized. The Protein A Cartridge is then washed to remove excess unbound proteins. The Desalting Cartridge is readied for use by reactivating with HEPES buffer. The Protein A Cartridge and Desalting Cartridge are then connected via the Luer lock fittings and the Elution Buffer is introduced. The eluate contains the purified IgG at a physiological pH. Both cartridges may be regenerated and stored for future use. The 10% nondenaturing sodium dodecyl sulphate polyacrylamide gel electrophoresis (SDS-PAGE) of sera and purified immunoglobulins was carried out [21].

Other immunoglobulins used in these experiments are the ostrich IgY purified by affinity chromatography [22] and commercially available IgGs from the pigs, mice, cats, rats, rabbits, and bovine IgG (Sigma-Aldrich).

**Figure 1:**
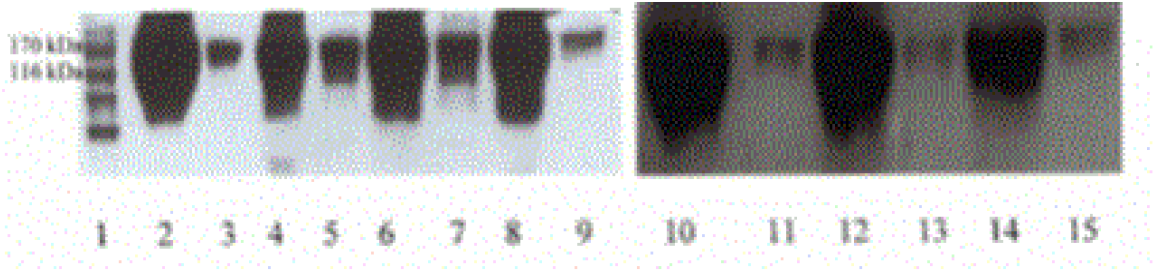
The 10% non-denaturing SDS-PAGE of sera and purified immunoglobulins (IgGs). Lane 1 molecular weight (MW) marker, lane 2 mule serum, lane 3 mule IgG, lane 4 donkey serum, lane 5 donkey IgG, lane 6 horse serum, lane 7 horse IgG, lane 8 dog serum, lane 9 dog IgG, lane 10 skunk serum, lane 11 skunk IgG, lane 12 coyote serum, lane 13 coyote IgG, lane 14 raccoon serum and lane 15 raccoon IgG. Purified IgGs have a molecular weight of approximately 150 Kda.

The chicken IgY fraction was isolated by the chloroform-polyethylene glycol (PEG) method [23]. The chicken egg was washed with warm water and the egg yolk was separated from the egg white. The membrane was broken, and the egg yolk collected and diluted 1:3 in phosphate buffered saline (PBS), pH 7.4. To 1/3 of the egg yolk mixture an equal volume of chloroform was added, the mixture was then shaken and centrifuged for 30 min (1000×g, RT). The supernatant was decanted and mixed with PEG 6000 (12%, w/v), stirred and incubated for 30 min (RT). The mixture was then centrifuged as previously described. The precipitate containing IgY was dissolved in PBS (pH 7.4) at a volume equivalent to 1/6 of the original volume of the egg yolk and dialyzed against 1 L of PBS (pH: 7.4 for 24 h at 4°C). The chicken IgY was removed from the dialysis tubing. IgY concentration was determined by the Bradford method. IgY samples were stored at – 20°C. Horseradish peroxidase (HRP) labelled SpL, SpA or SpG conjugates were prepared using the periodate method described by Nakane and Kawoi [24,25]. The PLAG-HRP conjugate along with SpLA-HRP, SpLG-HRP and SpAG-HRP were prepared by mixing at room temperature 50 μl of each SpL-HRP, SpA-HRP and SpG-HRP.

### Basic ELISA protocol

The 96 well microtitre plate is coated overnight at 4°C with 2 μg/μl per well of a mixture of SpA, SpG, SpL or their combinations SpAG, SpLG, SpLA or PLAG in carbonate-bicarbonate buffer pH 9.6. Then plate is treated with bovine serum albumin solution and washed 4X with PBS-Tween. 50 μl of immunoglobulins (1 mg/ml) is added and incubated for 1.30h at room temperature and the microplate is then rewashed 4X with PBS-Tween. Then 50 μl of a peroxidase-labeled-bacterial protein diluted to 1:3000-1:5000 in PBS-non-fat milk is added to each well and incubated for 1.30h at RT. After that, the plate is washed 4X with PBS-Tween. Pipette 50 μl of 3,3’,5,5’ - tetramethylbenzidine (TMB; Sigma-Aldrich) to each well. The reaction is stopped with 50 μl of 3M H2SO4 solution. The plate is visually assessed for the development of colour and read in a microplate reader at 450 nm. The assay is performed 3 times during the same day for the calculation of the intra-assay coefficient of variation, and once in 3 alternative days. A cut-off point is calculated as the mean of the optical density of negative controls times two [25].

The immunoglobulin-binding protein affinity to immunoglobulins are classified as weekly (+) and goes from the lower limit of the cut-off point to below the cut-off point times two value. Moderate binding affinity (++) goes from the lower limit of cut-off point times two to the below value of the cut-off point times 3 and strong binding affinity (+++) is an equal or higher value of the cut-off point times three. For instance, the mode of cut-off point calculated among the 35 assays was 0.30. The binding affinity of immunoglobulin binding-protein can be calculated as follows:

- From 0.30-0.59 (+)
- From 0.60-0.89 (++)
- From > 0.90 (+++)

#### Preparation of HIV immunogens

It was used a peptide fragment of gp120 from HIV gp120 (254-274): Cys-Thr-His-Gly-lle-Arg-Pro-Val-Val-Ser-Thr-Gln-LeuLeu-Leu-Asn-Gly-Ser-Leu-Ala-Glu [26]; that was conjugated to keyhole limpet hemocynin (KLH) by the glutaraldehyde method as previously described [27]. Chicken immunization with KLH-gp120 HIV fragment (254-274). Two healthy layer chickens (brown Leghorn), aged approximately 6 months, were injected intramuscularly at multiple sites on the breast with the peptide-KLH conjugate and they were immunized on day 0, 15, 60 as described previously. The eggs were collected postimmunization to test for anti-HIV antibodies [27].

### Oral immunization of cats

Feeding of cats with hyper-immune eggs The anti-gp120 positive eggs were fed to five (5) adult cats, 2-3 years old (1 male and 4 female). Each cat received on average, of 2 eggs diluted in 5 volumes of soya milk weekly for 10 weeks. Of the 5 cats used in the study, 3 were fed hyperimmune eggs (anti-gp120), while the remaining 2 cats were fed eggs from non-immunized chickens. And these two cats that were fed with eggs from non-immunized chickens were used as controls. Blood samples (2 ml) were collected from each cat, after completion of the feeding and were tested for anti-HIV antibodies by an indirect HIV ELISA [27].

Whole egg was fed to 3 rats (2-3 ml) by gastric intubation; weekly for 9 weeks and during the last week, they were fed daily. Two rats where fed HIV non-hyperimmune eggs as a negative control. Blood sample (0.5 ml) were drawn for testing for anti-HIV peptide antibodies by ELISA [27] and confirmative dot blot analyses [28].

### ELISA for detection of anti-HIV antibodies in eggs

The 96 well polystyrene microplates (U-shaped bottom) were coated with 50 ng of the synthetic HIVgp-120 peptide in coating buffer overnight at 4°C. The microplates were washed 4 times (PBSTween-20) and blocked with 3% non-fat milk in PBS, 25 μl/well, 1h, RT. The microplates were washed 4 times (PBS-Tween-20) and triplicates of 25 μl of WSF of the egg yolk in PBS non-fat milk were added. After incubation for 90 min at RT the microplates were washed 4 times and 25 μl of rabbit anti-chicken IgY-HRP diluted 1:30,000 was added and the microplates were then incubated for 1h at RT, washed four times. Tetramethylbenzidine (TMB) solution (50 μl) was added to each well. After a further incubation of 15 min in the dark, the reaction was stopped and read in a microplate reader at 450 nm. Geometric mean antibody titer was calculated using Perkins method as previously reported [27].

### Indirect ELISA for detection of anti-HIV antibodies in cats or rats

The ELISA assay described above (with modifications) was used to determine the presence of anti-HIV antibodies in cat or rat sera. The modifications were that triplicates of 1/16 dilutions of cat or rat serum samples in PBS non-fat milk were added. After incubation for 90 min at RT the microplates were washed 4X (PBS-Tween 20) and 25 μl of protein LA-HRP diluted 1:3000 (Sigma) was added and the microplates were then incubated for 1h at RT. Then, microplates were washed four times. Tetramethylbenzidine (TMB) solution (50 μl) was added to each well. After a further incubation of 15 min in the dark, the reaction was stopped and read with stop solution and the microplates were read in a microplate reader at 450 nm [27].

### Dot Blot analyses for the detection of anti-HIV peptide antibodies in cats and rats

A dot blot is a technique in molecular biology used to detect biomolecules, and for detecting, analyzing, and identifying proteins. Briefly, 2 μl of serum from cats, or rats in addition to human serum (positive control) and chicken IgY (a negative control) were blotted onto nitrocellulose paper placed in a BioDot SF apparatus (Bio-Rad Laboratories, Richmond, CA, USA). The membrane was blocked with 5 μL/well of fetal bovine serum with 1% Tris buffer saline, pH:7.4. After that, 5 μL of a commercial conjugate (peroxidase-labeled HIV proteins; Murex Diagnostics, Norcross, USA) was added to the nitrocellulose membrane for 1 hour at RT, washed and allowed to drain by gravity. Finally, 5 μL of the substrate 3,3’,5,5’-tetramethylbenzidine (Sigma-Aldrich) was added and the mixture was incubated for 20 min in the dark. The reaction was stopped by washing the wells with distilled water under a vacuum. The nitrocellulose membrane was left to dry, visualized and photographed [28].

### Statistical analysis

Statistical Package for the Social Sciences (SPSS) version 25 was used for the calculation of the coefficients of variation.

#### Ethical approval

This research was approved by the University of West Indies (UWI) Ethics Committee (Mona Campus) in Jamaica, West Indies.

## Results

### SpA-SpA sandwich ELISA

This ELISA is used to study the interaction of Staphylococcal protein-A (SpA) with diverse mammalian and avian immunoglobulins. The 96 well microtitre plate is coated overnight at 4°C with 2 μg/μl per well of SpA in carbonate-bicarbonate buffer pH 9.6. Then plate is treated with bovine serum albumin solution and washed 4X with PBS-Tween. 50 μl of immunoglobulins (1 mg/ml) is added and incubated for 1.30h at room temperature and the microplate is then rewashed 4X with PBS-Tween. Then 50 μl of peroxidase-labeled-SpA .2.0conjugate diluted 1:5000 in PBS-non-fat milk is added to each well and incubated for 1.30h at RT. The plate is washed 4X with PBS-Tween. Pipette 50 μl of 3,3’,5,5’ - tetramethylbenzidine (TMB; Sigma-Aldrich) to each well. The reaction is stopped with 50 μl of 3M H2SO4 solution. The plate is visually assessed for the development of colour and read in a microplate reader at 450 nm. A cut-off point is calculated as the mean of the optical density of negative controls times two. The higher the OD value the higher will be the affinity of SpA to avian immunoglobulins. The cut-off point was 0.30.

Table 1 shows the SpA-SpA sandwich ELISA. It was very effective. It demonstrated the strong binding affinity of IgG from diverse mammalian species including dog, skunk, raccoon, pig, mouse, rabbit, and cat. The chicken IgY did not bind to SpA. Some of these interactions have been previously cited [2,9,6,25].

**Table 1.** SpA-SpA sandwich ELISA.

| **Immunoglobulins** | **XOD results** | **Binding affinity** | **Coefficient of variation (CV) intra-assay (%)** | **CV inter-assay (%)** |
| --- | --- | --- | --- | --- |
| **Mule IgG** | **0.40** | **+** | **2.53** | **4.44** |
| **Donkey IgG** | **0.35** | **+** | **4.89** | **6.35** |
| **Horse IgG** | **0.56** | **++** | **3.67** | **5.38** |
| **Dog IgG** | **1.44** | **+++** | **3.03** | **4.71** |
| **Skunk IgG** | **1.10** | **+++** | **2.28** | **4.11** |
| **Coyote IgG** | **0.80** | **++** | **4.18** | **7.09** |
| **Raccoon IgG** | **1.52** | **+++** | **4.26** | **4.75** |
| **Ostrich IgY** | **0.42** | **+** | **3.30** | **9.68** |
| **Pig IgG** | **1.56** | **+++** | **4.14** | **7.47** |
| **Mouse IgG** | **1.48** | **+++** | **2.27** | **4.03** |
| **Bovine IgG** | **0.65** | **++** | **3.54** | **4.15** |
| **Rabbit IgG** | **1.28** | **+++** | **2.73** | **4.02** |
| **Chicken IgY** | **0.15** | **-** | **2.30** | **3.89** |
| **Cat IgG** | **1.46** | **+++** | **3.36** | **5.70** |
| **Rat IgG** | **0.39** | **+** | **4.09** | **6.83** |

### SpG-SpG Sandwich ELISA

This ELISA is used to study the interaction of streptococcal protein-G (SpG) with different mammalian and avian immunoglobulins. The 96 well microtitre plate is coated overnight at 4°C with 2 μg/μl per well of SpG in carbonate-bicarbonate buffer pH 9.6. Then plate is treated with bovine serum albumin solution and washed 4X with PBS-Tween. 50 μl of immunoglobulins (1 mg/ml) is added and incubated for 1.30h at room temperature and the microplate is then rewashed 4X with PBS-Tween. Then 50 μl of peroxidase-SpG conjugate diluted 1:5000 in PBS-non-fat milk is added to each well and incubated for 1.30h at RT. The plate is washed 4X with PBS-Tween. Pipette 50 μl of TMB (Sigma-Aldrich) to each well. The reaction is stopped with 50 μl of 3M H2SO4 solution. The plate is visually assessed for the development of colour and read in a microplate reader at 450 nm. A cut-off point is calculated as the mean of the optical density of negative controls times two. The higher the OD value the higher will be the affinity of SpG to mammalian immunoglobulins. The cut-off point is 0.28.

Table 2 shows the SpG-SpG sandwich ELISA, which depicted the interactions between SpG and IgGs from mules, donkeys, horses, dogs, skunks, pigs, mice, bovines, rats, and rabbits. None of the avian immunoglobulins bound to SpG. Some of these interactions were confirmatory results as horse, pig, bovine and rabbit reactivity [3,4].

**Table 2.** SpG-SpG Sandwich ELISA.

| **Immunoglobulins** | **XOD results** | **Binding affinity** | **Coefficient of variation (CV) intra-assay (%)** | **CV inter-assay (%)** |
| --- | --- | --- | --- | --- |
| **Mule IgG** | **0.96** | **+++** | **3.57** | **5.28** |
| **Donkey IgG** | **0.80** | **+++** | **3.89** | **5.35** |
| **Horse IgG** | **1.05** | **+++** | **4.98** | **7.38** |
| **Dog IgG** | **1.33** | **+++** | **3.03** | **8.70** |
| **Skunk IgG** | **1.24** | **+++** | **5.63** | **9.23** |
| **Coyote IgG** | **0.47** | **+** | **2.70** | **6.05** |
| **Raccoon IgG** | **0.42** | **+** | **3.28** | **6.60** |
| **Ostrich IgY** | **0.08** | **-** | **2.85** | **8.68** |
| **Pig IgG** | **1.36** | **+++** | **5.09** | **10.3** |
| **Mouse IgG** | **1.55** | **+++** | **3.06** | **7.00** |
| **Bovine IgG** | **1.49** | **+++** | **4.96** | **8.63** |
| **Rabbit IgG** | **1.25** | **+++** | **5.83** | **9.28** |
| **Chicken IgY** | **0.14** | **-** | **3.50** | **5.06** |
| **Cat IgG** | **0.18** | **-** | **3.01** | **4.60** |
| **Rat IgG** | **0.93** | **+++** | **5.83** | **8.74** |

### SpL-SpL Sandwich ELISA

This ELISA is used to study the interaction of a protein-L (SpL) with different immunoglobulin preparations from various mammalian and avian species. The 96 well microtiter plate is coated overnight at 4°C with 2 μg/μl per well of SpL in carbonate-bicarbonate buffer pH 9.6. Then plate is treated with bovine serum albumin solution and washed 4X with PBS-Tween. 50 μl of immunoglobulins (1 mg/ml) is added and incubated for 1h at room temperature and the microplate is rewashed 4X with PBS-Tween. Then, 50 μl of peroxidase-labeled SpL conjugate diluted 1:5000 in PBS-non-fat milk is added to each well and incubated for 1h at RT. The plate is washed 4X with PBS-Tween. 50 μl of 4 mg/ml o-phenylenediamine solution (OPD) is added and the plate is incubated 15 minutes at RT in the dark. The reaction is stopped with 50 μl of 3M H2SO4 solution. The plate is visually assessed for the development of colour and read in a microplate reader at 492 nm. A cut-off point is calculated as the mean of the optical density of negative controls times two. The higher the OD value the higher will be the affinity of SpL to immunoglobulins. The cut-off point is 0.28.

Table 3 shows SpL that interacts with fewer immunoglobulins than its counterpart SpA and SpG. IgGs from pigs and mice reacted strongly with SpL. Other species as dog, raccoon and rabbit reacted weakly [5,7].

**Table 3.** SpL-SpL Sandwich ELISA.

| Immunoglobulins | XOD results | Binding affinity | Coefficient of variation (CV) intra-assay (%) | CV inter-assay (%) |
| --- | --- | --- | --- | --- |
| Mule IgG | 0.08 | - | 2.97 | 3.56 |
| Donkey IgG | 0.07 | - | 2.50 | 4.09 |
| Horse IgG | 0.1 | - | 2.65 | 3.17 |
| Dog IgG | 0.39 | + | 4.12 | 5.65 |
| Skunk IgG | 0.13 | - | 2.46 | 5.10 |
| Coyote IgG | 0.09 | - | 3.45 | 5.15 |
| Raccoon IgG | 0.41 | + | 4.04 | 6.89 |
| Ostrich IgY | 0.15 | - | 2.69 | 5.50 |
| Pig IgG | 0.98 | +++ | 3.87 | 6.37 |
| Mouse IgG | 1.07 | +++ | 2.48 | 4.98 |
| Bovine IgG | 0.07 | - | 4.07 | 6.05 |
| Rabbit IgG | 0.38 | + | 3.40 | 6.61 |
| Chicken IgY | 0.14 | - | 2.61 | 4.05 |
| Cat IgG | 0.11 | - | 3.06 | 4.98 |
| Rat IgG | 0.09 | - | 2.78 | 3.95 |

### SpLG-SpLG Sandwich ELISA

This ELISA is used to study the interaction of a protein-LG (SpLG) with different immunoglobulin preparations from mammalian and avian species. The 96 well microtitre plate is coated overnight at 4°C with 2 μg/μl per well of a mixture of protein-L and protein-G in carbonate-bicarbonate buffer pH 9.6. Then plate is treated with bovine serum albumin solution and washed 4X with PBS-Tween. 50 μl of immunoglobulins (1 mg/ml) is added and incubated for 1h at room temperature and the microplate is rewashed 4X with PBS-Tween. Then 50 μl of peroxidase-labeled SpLG conjugate diluted 1:5000 in PBS-non-fat milk is added to each well and incubated for 1h at RT. The plate is washed 4X with PBS-Tween. 50 μl of 4 mg/ml o-phenylenediamine solution (OPD) is added and the plate is incubated 15 minutes at RT in the dark. The reaction is stopped with 50 μl of 3M H2SO4 solution. The plate is visually assessed for the development of colour and read in a microplate reader at 492 nm. A cut-off point is calculated as the mean of the optical density of negative controls times two. The higher the OD value the higher will be the affinity of SpLG to molecules. The cut-off is 0.32.

The SpLG conjugate used for the standardization of this immunoassay was prepared by mixing up SpL-HRP and SpG-HRP. It combines the binding affinities of both proteins. Table 4 depicts this assay, where IgGs from ten different species interacted strongly with SpLG. It did not react with cat IgG and chicken IgY but reacted weakly with IgGs from coyotes and raccoons.

**Table 4.**
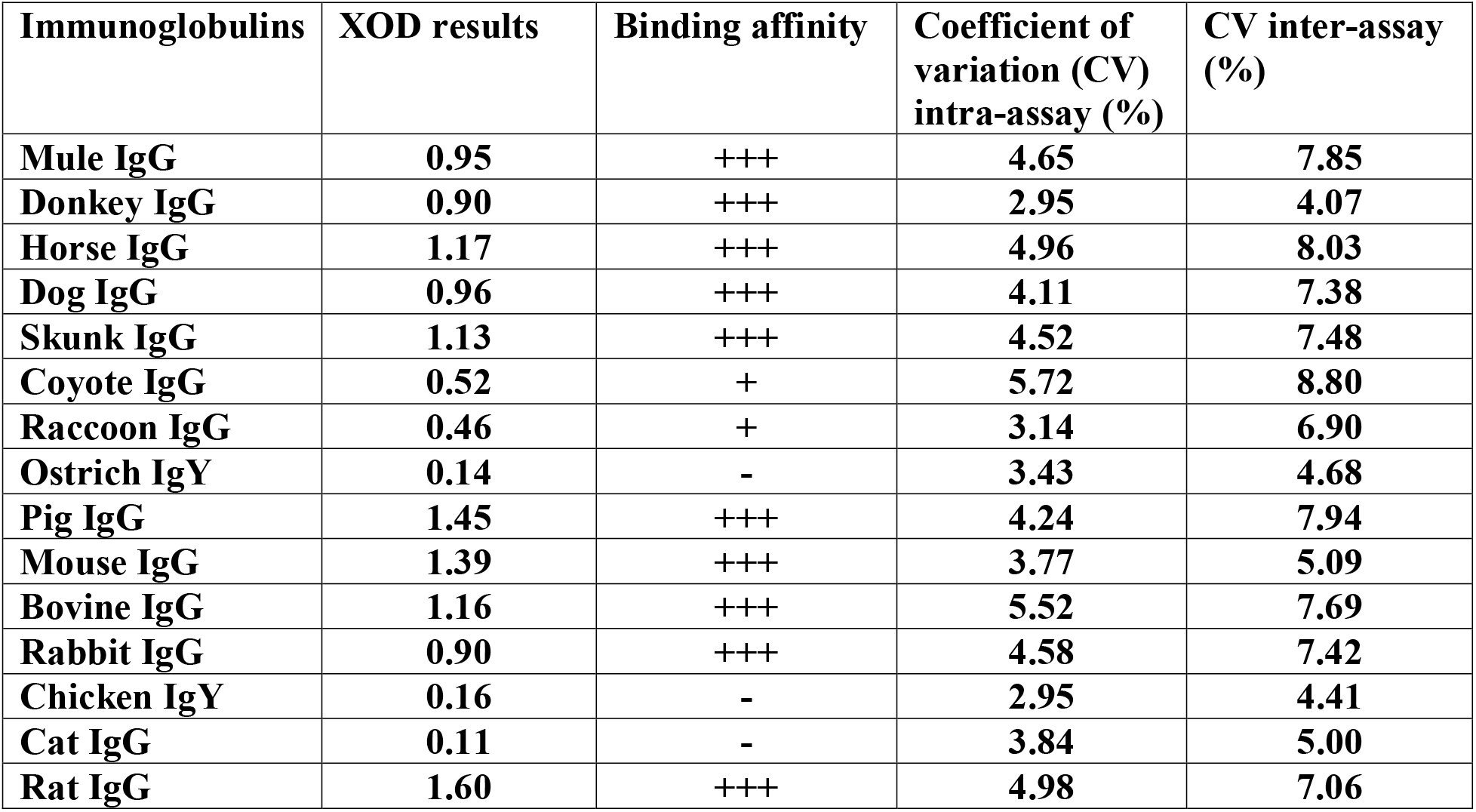
SpLG-SpLG Sandwich ELISA.

| **Immunoglobulins** | **XOD results** | **Binding affinity** | **Coefficient of variation (CV) intra-assay (%)** | **CV inter-assay (%)** |
| --- | --- | --- | --- | --- |
| **Mule IgG** | **0.95** | **+++** | **4.65** | **7.85** |
| **Donkey IgG** | **0.90** | **+++** | **2.95** | **4.07** |
| **Horse IgG** | **1.17** | **+++** | **4.96** | **8.03** |
| **Dog IgG** | **0.96** | **+++** | **4.11** | **7.38** |
| **Skunk IgG** | **1.13** | **+++** | **4.52** | **7.48** |
| **Coyote IgG** | **0.52** | **+** | **5.72** | **8.80** |
| **Raccoon IgG** | **0.46** | **+** | **3.14** | **6.90** |
| **Ostrich IgY** | **0.14** | **-** | **3.43** | **4.68** |
| **Pig IgG** | **1.45** | **+++** | **4.24** | **7.94** |
| **Mouse IgG** | **1.39** | **+++** | **3.77** | **5.09** |
| **Bovine IgG** | **1.16** | **+++** | **5.52** | **7.69** |
| **Rabbit IgG** | **0.90** | **+++** | **4.58** | **7.42** |
| **Chicken IgY** | **0.16** | **-** | **2.95** | **4.41** |
| **Cat IgG** | **0.11** | **-** | **3.84** | **5.00** |
| **Rat IgG** | **1.60** | **+++** | **4.98** | **7.06** |

### SpLA-SpLA Sandwich ELISA

This ELISA is used to study the interaction of a protein-LA (SpLA) with different immunoglobulin preparations from mammalian and avian species. The 96 well microtitre plate is coated overnight at 4°C with 2 μg/μl per well of a mixture of protein-L and protein-A in carbonate-bicarbonate buffer pH 9.6. Then plate is treated with bovine serum albumin solution and washed 4X with PBS-Tween. 50 μl of immunoglobulins (1 mg/ml) is added and incubated for 1h at room temperature and the microplate is rewashed 4X with PBS-Tween. Then 50 μl of peroxidase-labeled SpLA conjugate diluted 1:5000 in PBS-non-fat milk is added to each well and incubated for 1h at RT. The plate is washed 4X with PBS-Tween. 50 μl of 4 mg/ml o-phenylenediamine solution (OPD) is added and the plate is incubated 15 minutes at RT in the dark. The reaction is stopped with 50 μl of 3M H2SO4 solution. The plate is visually assessed for the development of colour and read in a microplate reader at 492 nm. A cut-off point is calculated as the mean of the optical density of negative controls times two. The higher the OD value the higher will be the affinity of SpLA to mammalian immunoglobulins. The cut-off point is 0.30.

Table 5 depicts SpLA interacting strongly with a range of immunoglobulins. Very high affinity was detected between SpLA and IgGs from pigs, mice, and other few species.

**Table 5.** SpLA-SpLA Sandwich ELISA.

| **Immunoglobulins** | **XOD results** | **Binding affinity** | **Coefficient of variation (CV) intra-assay (%)** | **CV inter-assay (%)** |
| --- | --- | --- | --- | --- |
| **Mule IgG** | **0.44** | **+** | **3.14** | **5.46** |
| **Donkey IgG** | **0.50** | **+** | **3.36** | **5.78** |
| **Horse IgG** | **0.75** | **++** | **3.38** | **6.04** |
| **Dog IgG** | **1.05** | **+++** | **5.13** | **7.43** |
| **Skunk IgG** | **1.01** | **+++** | **3.50** | **6.41** |
| **Coyote IgG** | **0.80** | **++** | **4.25** | **6.36** |
| **Raccoon IgG** | **1.25** | **+++** | **4.77** | **8.29** |
| **Ostrich IgY** | **0.45** | **+** | **3.05** | **5.67** |
| **Pig IgG** | **1.46** | **+++** | **3.88** | **6.11** |
| **Mouse IgG** | **1.50** | **+++** | **4.08** | **6.95** |
| **Bovine IgG** | **0.52** | **+** | **3.62** | **5.45** |
| **Rabbit IgG** | **1.05** | **+++** | **4.27** | **6.80** |
| **Chicken IgY** | **0.15** | **-** | **3.30** | **4.04** |
| **Cat IgG** | **1.05** | **+++** | **3.75** | **6.02** |
| **Rat IgG** | **0.50** | **+** | **2.96** | **4.63** |

### SpAG-SpAG Sandwich ELISA

This ELISA is used to study the interaction of protein-AG (SpAG) with diverse immunoglobulins. The 96 well microtitre plate is coated overnight at 4°C with 2 μg/μl per well of a mixture of SpA with SpG in carbonate-bicarbonate buffer pH 9.6. Then plate is treated with bovine serum albumin solution and washed 4X with PBS-Tween. 50 μl of immunoglobulins (1 mg/ml) is added and incubated for 1.30h at room temperature and the microplate is then rewashed 4X with PBS-Tween. Then 50 μl of peroxidase-labeled-SpAG conjugate diluted 1:5000 in PBS-non-fat milk is added to each well and incubated for 1.30h at RT. After that, the plate is washed 4X with PBS-Tween. Pipette 50 μl of 3,3’,5,5’ - tetramethylbenzidine (TMB; Sigma-Aldrich) to each well. The reaction is stopped with 50 μl of 3M H2SO4 solution. The plate is visually assessed for the development of colour and read in a microplate reader at 450 nm. A cut-off point is calculated as the mean of the optical density of negative controls times two. The higher the OD value the higher will be the binding affinity of SpAG to immunoglobulins. The cut-off point was 0.30.

Table 6 shows that SpAG is a versatile protein, which reacted strongly with 13 out of 15 IgGs. It reacted strongly with the 86.6% of the Ig panel.

**Table 6.** SpAG-SpAG Sandwich ELISA.

| **Immunoglobulins** | **XOD results** | **Binding affinity** | **Coefficient of variation (CV) intra-assay (%)** | **CV inter-assay (%)** |
| --- | --- | --- | --- | --- |
| **Mule IgG** | **0.92** | **+++** | **3.78** | **6.74** |
| **Donkey IgG** | **1.05** | **+++** | **2.64** | **5.30** |
| **Horse IgG** | **0.95** | **+++** | **4.15** | **7.85** |
| **Dog IgG** | **1.26** | **+++** | **3.94** | **5.71** |
| **Skunk IgG** | **1.24** | **+++** | **3.29** | **4.78** |
| **Coyote IgG** | **0.99** | **+++** | **3.88** | **6.67** |
| **Raccoon IgG** | **0.95** | **+++** | **5.16** | **8.77** |
| **Ostrich IgY** | **0.45** | **+** | **2.42** | **3.21** |
| **Pig IgG** | **1.43** | **+++** | **3.89** | **6.60** |
| **Mouse IgG** | **1.41** | **+++** | **2.67** | **4.79** |
| **Bovine IgG** | **1.14** | **+++** | **3.35** | **5.16** |
| **Rabbit IgG** | **1.33** | **+++** | **4.38** | **6.84** |
| **Chicken IgY** | **0.15** | **-** | **2.96** | **4.43** |
| **Cat IgG** | **1.34** | **+++** | **3.17** | **4.83** |
| **Rat IgG** | **1.10** | **+++** | **2.89** | **5.01** |

### SpA and SpLA sandwich ELISA

This ELISA is used to study the interactions between Staphylococcal protein-A (SpA) and protein-LA (SpLA) with different immunoglobulin preparations from mammalian and avian species. The 96 well microtitre plate is coated overnight at 4°C with 2 μg/μl per well of SpA in carbonate-bicarbonate buffer pH 9.6. Then plate is treated with bovine serum albumin solution and washed 4X with PBS-Tween. 50 μl of immunoglobulins (1 mg/ml) is added and incubated for 1h at room temperature and the microplate is rewashed 4X with PBS-Tween. Then 50 μl of peroxidase-labeled SpLA conjugate diluted 1:5000 in PBS-non-fat milk is added to each well and incubated for 1h at RT. The plate is washed 4X with PBS-Tween. 50 μl of 4 mg/ml o-phenylenediamine solution (OPD) is added and the plate is incubated 15 minutes at RT in the dark. The reaction is stopped with 50 μl of 3M H2SO4 solution. The plate is visually assessed for the development of colour and read in a microplate reader at 492 nm. The cut-off point is 0.32.

Table 7 depicts a hybrid ELISA that demonstrates the binding affinity of both SpA and SpLA. IgG in this ELISA are more likely to interact with SpA. As demonstrated in previous ELISA the binding affinity of SpA surpasses that of SpL. IgGs from dogs, skunks, pigs, mice, rabbits, and cats reacted strongly with SpA. But pig and mouse immunoglobulins had the highest binding affinities since they are capable of strongly reacting to both SpA and SpL. There is a report of the creation of a versatile hybrid recombinant SpLA [29].

**Table 7.** SpA and SpLA sandwich ELISA.

| **Immunoglobulins** | **XOD results** | **Binding affinity** | **Coefficient of variation (CV) intra-assay (%)** | **CV inter-assay (%)** |
| --- | --- | --- | --- | --- |
| **Mule IgG** | **0.45** | **+** | **2.53** | **4.44** |
| **Donkey IgG** | **0.55** | **+** | **4.89** | **6.35** |
| **Horse IgG** | **0.68** | **++** | **3.67** | **5.38** |
| **Dog IgG** | **1.40** | **+++** | **3.03** | **4.71** |
| **Skunk IgG** | **1.21** | **+++** | **2.28** | **4.11** |
| **Coyote IgG** | **0.75** | **++** | **4.18** | **7.09** |
| **Raccoon IgG** | **1.47** | **+++** | **4.26** | **4.75** |
| **Ostrich IgY** | **0.53** | **+** | **3.30** | **9.68** |
| **Pig IgG** | **1.50** | **+++** | **4.14** | **7.47** |
| **Mouse IgG** | **1.59** | **+++** | **2.27** | **4.03** |
| **Bovine IgG** | **0.70** | **++** | **3.54** | **4.15** |
| **Rabbit IgG** | **1.05** | **+++** | **2.73** | **4.02** |
| **Chicken IgY** | **0.16** | **-** | **2.30** | **3.89** |
| **Cat IgG** | **1.45** | **+++** | **3.36** | **5.70** |
| **Rat IgG** | **0.53** | **+** | **4.09** | **6.83** |

### SpA and SpLG sandwich ELISA

This ELISA is used to study the interactions between Staphylococcal protein-A (SpA) and protein-LG (SpLG) with different immunoglobulin preparations from mammalian and avian species. The 96 well microtitre plate is coated overnight at 4°C with 2 μg/μl per well of SpA in carbonate-bicarbonate buffer pH 9.6. Then plate is treated with bovine serum albumin solution and washed 4X with PBS-Tween. 50 μl of immunoglobulins (1 mg/ml) is added and incubated for 1h at room temperature and the microplate is rewashed 4X with PBS-Tween. Then 50 μl of peroxidase-labeled SpLG conjugate diluted 1:5000 in PBS-non-fat milk is added to each well and incubated for 1h at RT. The plate is washed 4X with PBS-Tween. 50 μl of 4 mg/ml o-phenylenediamine solution (OPD) is added and the plate is incubated 15 minutes at RT in the dark. The reaction is stopped with 50 μl of 3M H2SO4 solution. The plate is visually assessed for the development of colour and read in a microplate reader at 492 nm. A cut-off point is calculated as the mean of the optical density of negative controls times two. The cut-off point was 0.32.

Table 8 shows a type of immunoassay, where immunoglobulins serve as a bridge between two different bacterial proteins. SpA and SpLG react strongly with equine Igs, and IgGs of dogs, skunks, raccoons, pigs, mice, rabbits, and rats. It did not react with chicken IgY, which in fact is the negative control used to calculate the cut-off point.

**Table 8.** SpA and SpLG sandwich ELISA.

| **Immunoglobulins** | **XOD results** | **Binding affinity** | **Coefficient of variation (CV) intra-assay (%)** | **CV inter-assay (%)** |
| --- | --- | --- | --- | --- |
| **Mule IgG** | **1.25** | **+++** | **3.67** | **6.95** |
| **Donkey IgG** | **1.02** | **+++** | **4.08** | **8.05** |
| **Horse IgG** | **1.37** | **+++** | **3.85** | **5.24** |
| **Dog IgG** | **1.45** | **+++** | **3.95** | **6.00** |
| **Skunk IgG** | **1.23** | **+++** | **4.74** | **6.25** |
| **Coyote IgG** | **0.47** | **+** | **3.89** | **5.97** |
| **Raccoon IgG** | **1.30** | **+++** | **2.96** | **4.75** |
| **Ostrich IgY** | **0.52** | **+** | **4.55** | **5.90** |
| **Pig IgG** | **1.05** | **+++** | **3.06** | **6.66** |
| **Mouse IgG** | **1.37** | **+++** | **4.50** | **7.57** |
| **Bovine IgG** | **0.70** | **++** | **6.01** | **9.28** |
| **Rabbit IgG** | **1.15** | **+++** | **4.07** | **6.37** |
| **Chicken IgY** | **0.16** | **-** | **4.98** | **8.05** |
| **Cat IgG** | **0.46** | **+** | **4.82** | **7.42** |
| **Rat IgG** | **1.40** | **+++** | **5.86** | **9.23** |

### Table 9. SpA and SpAG sandwich ELISA

This ELISA is used to study the interactions between Staphylococcal protein-A (SpA) and protein-AG (SpAG) with different immunoglobulin preparations from mammalian and avian species. The 96 well microtitre plate is coated overnight at 4°C with 2 μg/μl per well of SpA in carbonate-bicarbonate buffer pH 9.6. Then plate is treated with bovine serum albumin solution and washed 4X with PBS-Tween. 50 μl of immunoglobulins (1 mg/ml) is added and incubated for 1h at room temperature and the microplate is rewashed 4X with PBS-Tween. Then 50 μl of peroxidase-labeled SpAG conjugate diluted 1:5000 in PBS-non-fat milk is added to each well and incubated for 1h at RT. The plate is washed 4X with PBS-Tween. 50 μl of 4 mg/ml o-phenylenediamine solution is added and the plate is incubated 15 minutes at RT in the dark. The reaction is stopped with 50 μl of 3M H2SO4 solution. The plate is visually assessed for the development of colour and read in a microplate reader at 492 nm. A cut-off point is calculated as the mean of the optical density of negative controls times two. The cut-off point is 0.34.

Table 9 depicts immunoglobulins reacting strongly with SpA and SpAG. In this assay, except ostrich IgY and IgGs from rats interacted with the immunoglobulin-binding protein.

SpA and SpAG sandwich ELISA.

SpA and SpAG sandwich ELISA
| Immunoglobulins | XOD results | Binding affinity | Coefficient of variation (CV) intra-assay (%) | CV inter-assay (%) |
| --- | --- | --- | --- | --- |
| Mule IgG | 1.18 | +++ | 4.18 | 7.28 |
| Donkey IgG | 1.26 | +++ | 3.78 | 5.95 |
| Horse IgG | 1.42 | +++ | 3.06 | 5.70 |
| Dog IgG | 1.60 | +++ | 2.79 | 4.86 |
| Skunk IgG | 1.33 | +++ | 4.05 | 6.37 |
| Coyote IgG | 1.20 | +++ | 3.80 | 6.21 |
| Raccoon IgG | 1.15 | +++ | 3.68 | 5.07 |
| Ostrich IgY | 0.49 | + | 4.86 | 7.10 |
| Pig IgG | 1.67 | +++ | 2.96 | 4.97 |
| Mouse IgG | 1.57 | +++ | 4.00 | 7.67 |
| Bovine IgG | 1.08 | +++ | 4.29 | 7.42 |
| Rabbit IgG | 1.24 | +++ | 4.76 | 6.60 |
| Chicken IgY | 0.17 | - | 3.50 | 5.73 |
| Cat IgG | 1.13 | +++ | 3.79 | 6.01 |
| Rat IgG | 0.48 | + | 4.86 | 6.52 |

### SpG and SpLA sandwich ELISA

This ELISA is used to study the interactions between Staphylococcal protein-G (SpG) and protein-LA (SpLA) with different immunoglobulin preparations from mammalian and avian species. The 96 well microtitre plate is coated overnight at 4°C with 2 μg/μl per well of SpG in carbonate-bicarbonate buffer pH 9.6. Then plate is treated with bovine serum albumin solution and washed 4X with PBS-Tween. 50 μl of immunoglobulins (1 mg/ml) is added and incubated for 1h at room temperature and the microplate is rewashed 4X with PBS-Tween. Then 50 μl of peroxidase-labeled SpLA conjugate diluted 1:5000 in PBS-non-fat milk is added to each well and incubated for 1h at RT. The plate is washed 4X with PBS-Tween. 50 μl of 4 mg/ml o-phenylenediamine solution (OPD) is added and the plate is incubated 15 minutes at RT in the dark. The reaction is stopped with 50 μl of 3M H2SO4 solution. The plate is visually assessed for the development of colour and read in a microplate reader at 492 nm. A cut-off point is calculated as the mean of the optical density of negative controls times two. The cut-off point is 0.36.

In this immunoassay both SpG and SpLA react with the entire panel of immunoglobulins as shown in Table 10. All IgGs react effectively with both bacterial protein except cat and rat IgGs that reacted weakly.

**Table 10.** SpG and SpLA sandwich ELISA.

| **Immunoglobulins** | **XOD results** | **Binding affinity** | **Coefficient of variation (CV) intra-assay (%)** | **CV inter-assay (%)<sup>78</sup>** |
| --- | --- | --- | --- | --- |
| **Mule IgG** | **1.15** | **+++** | **3.27** | **7.28** |
| **Donkey IgG** | **1.24** | **+++** | **3.86** | **6.04** |
| **Horse IgG** | **1.17** | **+++** | **3.56** | **5.86** |
| **Dog IgG** | **1.48** | **+++** | **3.60** | **5.80** |
| **Skunk IgG** | **1.20** | **+++** | **3.09** | **5.94** |
| **Coyote IgG** | **0.93** | **+++** | **4.86** | **7.04** |
| **Raccoon IgG** | **1.04** | **+++** | **4.06** | **7.96** |
| **Ostrich IgY** | **0.65** | **++** | **3.75** | **6.87** |
| **Pig IgG** | **1.55** | **+++** | **3.75** | **5.80** |
| **Mouse IgG** | **1.68** | **+++** | **4.38** | **6.03** |
| **Bovine IgG** | **0.70** | **++** | **4.47** | **6.78** |
| **Rabbit IgG** | **1.32** | **+++** | **3.21** | **5.27** |
| **Chicken IgY** | **0.18** | **-** | **4.81** | **8.08** |
| **Cat IgG** | **0.37** | **+** | **3.44** | **6.80** |
| **Rat IgG** | **0.52** | **+** | **3.86** | **5.11** |

### SpG and SpLG sandwich ELISA

This ELISA is used to study the interactions between Staphylococcal protein-G (SpG) and protein-LG (SpLG) with different immunoglobulin preparations from mammalian and avian species. The 96 well microtitre plate is coated overnight at 4°C with 2 μg/μl per well of SpG in carbonate-bicarbonate buffer pH 9.6. Then plate is treated with bovine serum albumin solution and washed 4X with PBS-Tween. 50 μl of immunoglobulins (1 mg/ml) is added and incubated for 1h at room temperature and the microplate is rewashed 4X with PBS-Tween. Then 50 μl of peroxidase-labeled SpLG conjugate diluted 1:5000 in PBS-non-fat milk is added to each well and incubated for 1h at RT. The plate is washed 4X with PBS-Tween. 50 μl of 4 mg/ml o-phenylenediamine solution (OPD) is added and the plate is incubated 15 minutes at RT in the dark. The reaction is stopped with 50 μl of 3M H2SO4 solution. The plate is visually assessed for the development of colour and read in a microplate reader at 492 nm. A cut-off point is calculated as the mean of the optical density of negative controls times two. The cut-off point is 0.36.

In the SpG-SpLG ELISA, IgGs from equines, dogs, skunks, pigs, mice, bovines, and rabbits strongly bound to both bacterial proteins as shown in Table 11. However, cat IgG was below the cut-off point.

**Table 11.** SpG and SpLG sandwich ELISA.

| Immunoglobulins | XOD results | Binding affinity | Coefficient of variation (CV) intra-assay (%) | CV inter-assay (%) |
| --- | --- | --- | --- | --- |
| Mule IgG | 1.25 | +++ | 2.68 | 5.08 |
| Donkey IgG | 1.14 | +++ | 2.95 | 4.86 |
| Horse IgG | 1.30 | +++ | 3.15 | 5.76 |
| Dog IgG | 1.53 | +++ | 2.69 | 5.06 |
| Skunk IgG | 1.13 | +++ | 3.09 | 6.20 |
| Coyote IgG | 0.75 | ++ | 2.96 | 4.16 |
| Raccoon IgG | 0.42 | + | 4.52 | 8.15 |
| Ostrich IgY | 0.51 | + | 3.89 | 7.02 |
| Pig IgG | 1.38 | +++ | 4.34 | 6.70 |
| Mouse IgG | 1.35 | +++ | 3.66 | 5.81 |
| Bovine IgG | 1.05 | +++ | 4.01 | 6.22 |
| Rabbit IgG | 1.42 | +++ | 5.15 | 7.88 |
| Chicken IgY | 0.18 | - | 2.96 | 4.28 |
| Cat IgG | 0.25 | - | 4.75 | 8.10 |
| Rat IgG | 0.75 | ++ | 2.97 | 5.09 |

### SpG and SpAG sandwich ELISA

This ELISA is used to study the interactions between Staphylococcal protein-G (SpG) and protein-AG (SpAG) with different immunoglobulin preparations from mammalian and avian species. The 96 well microtitre plate is coated overnight at 4°C with 2 μg/μl per well of SpG in carbonate-bicarbonate buffer pH 9.6. Then plate is treated with bovine serum albumin solution and washed 4X with PBS-Tween. 50 μl of immunoglobulins (1 mg/ml) is added and incubated for 1h at room temperature and the microplate is rewashed 4X with PBS-Tween. Then 50 μl of peroxidase-labeled SpAG conjugate diluted 1:5000 in PBS-non-fat milk is added to each well and incubated for 1h at RT. The plate is washed 4X with PBS-Tween. 50 μl of 4 mg/ml o-phenylenediamine solution (OPD) is added and the plate is incubated 15 minutes at RT in the dark. The reaction is stopped with 50 μl of 3M H2SO4 solution. The plate is visually assessed for the development of colour and read in a microplate reader at 492 nm. A cut-off point is calculated as the mean of the optical density of negative controls times two. The cut-off point is 0.32.

The cat IgG binds weakly to SpG. Some authors have shown no reactivity at all [6]. However, the panel of 35 ELISAs in this study assure a 1+ binding between cat IgG and SpG. On the other hand, SpG and SpAG strongly interact with IgGs from much species including mule, donkey, dog, skunk, pig, and others as shown in Table 12.

**Table 12.** SpG and SpAG sandwich ELISA.

| **Immunoglobulins** | **XOD results** | **Binding affinity** | **Coefficient of variation (CV) intra-assay (%)** | **CV inter-assay (%)** |
| --- | --- | --- | --- | --- |
| **Mule IgG** | **0.98** | **+++** | **4.13** | **7.58** |
| **Donkey IgG** | **1.03** | **+++** | **3.60** | **6.26** |
| **Horse IgG** | **1.25** | **+++** | **4.15** | **7.05** |
| **Dog IgG** | **1.31** | **+++** | **3.22** | **5.46** |
| **Skunk IgG** | **1.37** | **+++** | **4.14** | **7.05** |
| **Coyote IgG** | **0.75** | **++** | **2.89** | **5.27** |
| **Raccoon IgG** | **0.87** | **++** | **3.65** | **5.08** |
| **Ostrich IgY** | **0.52** | **+** | **3.49** | **5.75** |
| **Pig IgG** | **1.35** | **+++** | **4.06** | **6.35** |
| **Mouse IgG** | **1.40** | **+++** | **4.18** | **7.25** |
| **Bovine IgG** | **1.17** | **+++** | **3.79** | **7.17** |
| **Rabbit IgG** | **1.23** | **+++** | **3.88** | **6.70** |
| **Chicken IgY** | **0.16** | **-** | **3.05** | **5.12** |
| **Cat IgG** | **0.36** | **+** | **4.85** | **8.19** |
| **Rat IgG** | **0.70** | **++** | **3.45** | **6.67** |

### SpL and SpAG sandwich ELISA

This ELISA is used to study the interactions between Staphylococcal protein-L (SpL) and protein-AG with different immunoglobulin preparations from mammalian and avian species. The 96 well microtitre plate is coated overnight at 4°C with 2 μg/μl per well of SpL in carbonate-bicarbonate buffer pH 9.6. Then plate is treated with bovine serum albumin solution and washed 4X with PBS-Tween. 50 μl of immunoglobulins (1 mg/ml) is added and incubated for 1h at room temperature and the microplate is rewashed 4X with PBS-Tween. Then, 50 μl of peroxidase-labeled SpAG conjugate diluted 1:5000 in PBS-non-fat milk is added to each well and incubated for 1h at RT. The plate is washed 4X with PBS-Tween. 50 μl of 4 mg/ml o-phenylenediamine solution (OPD) is added and the plate is incubated 15 minutes at RT in the dark. The reaction is stopped with 50 μl of 3M H2SO4 solution. The plate is visually assessed for the development of colour and read in a microplate reader at 492 nm. A cut-off point is calculated as the mean of the optical density of negative controls times two. The cut-off point was 0.28.

In this ELISA depicted in Table 13, the 3 bacterial proteins have the capacity to react strongly with IgGs from several species including pig, skunk, and dog. Immunoglobulins from ten different animal species were below the cut-off point.

**Table 13.** SpL and SpAG sandwich ELISA.

| Immunoglobulins | XOD results | Binding affinity | Coefficient of variation (CV) intra-assay (%) | CV inter-assay (%) |
| --- | --- | --- | --- | --- |
| Mule IgG | 0.1 | - | 2.85 | 4.50 |
| Donkey IgG | 0.08 | - | 2.46 | 3.07 |
| Horse IgG | 0.11 | - | 3.15 | 4.84 |
| Dog IgG | 0.41 | + | 4.12 | 5.61 |
| Skunk IgG | 0.11 | - | 3.46 | 4.53 |
| Coyote IgG | 0.09 | - | 2.45 | 3.94 |
| Raccoon IgG | 0.42 | + | 3.46 | 5.83 |
| Ostrich IgY | 0.18 | - | 2.96 | 4.40 |
| Pig IgG | 1.15 | +++ | 3.80 | 5.06 |
| Mouse IgG | 1.12 | +++ | 2.48 | 3.98 |
| Bovine IgG | 0.09 | - | 2.77 | 5.57 |
| Rabbit IgG | 0.34 | + | 3.43 | 5.60 |
| Chicken IgY | 0.14 | - | 2.10 | 4.56 |
| Cat IgG | 0.10 | - | 2.51 | 3.90 |
| Rat IgG | 0.01 | - | 3.01 | 4.96 |

### SpL and SpLG sandwich ELISA

This ELISA is used to study the interactions between Staphylococcal protein-L (SpL) and protein-LG with different immunoglobulin preparations from mammalian and avian species. The 96 well microtitre plate is coated overnight at 4°C with 2 μg/μl per well of SpL in carbonate-bicarbonate buffer pH 9.6. Then plate is treated with bovine serum albumin solution and washed 4X with PBS-Tween. 50 μl of immunoglobulins (1 mg/ml) is added and incubated for 1h at room temperature and the microplate is rewashed 4X with PBS-Tween. Then, 50 μl of peroxidase-labeled SpLG conjugate diluted 1:5000 in PBS-non-fat milk is added to each well and incubated for 1h at RT. The plate is washed 4X with PBS-Tween. 50 μl of 4 mg/ml o-phenylenediamine solution (OPD) is added and the plate is incubated 15 minutes at RT in the dark. The reaction is stopped with 50 μl of 3M H2SO4 solution. The plate is visually assessed for the development of colour and read in a microplate reader at 492 nm. A cut-off point is calculated as the mean of the optical density of negative controls times two. The cut-off point was 0.32.

In this immunoassay only some species bind to the solid phase since the binding of SpL to immunoglobulins is limited as compared of that of SpA and SpG. AS shown in Table 14 only pig and mouse IgGs interact strongly with the bacterial receptors (SpL and SpLG).

**Table 14.** SpL and SpLG sandwich ELISA.

| **Immunoglobulins** | **XOD results** | **Binding affinity** | **Coefficient of variation (CV) intra-assay (%)** | **CV inter-assay (%)** |
| --- | --- | --- | --- | --- |
| **Mule IgG** | **0.15** | - | **3.85** | **5.50** |
| **Donkey IgG** | **0.12** | - | **3.46** | **4.00** |
| **Horse IgG** | **0.17** | - | **2.25** | **3.73** |
| **Dog IgG** | **0.38** | + | **3.30** | **4.68** |
| **Skunk IgG** | **0.20** | - | **4.01** | **7.55** |
| **Coyote IgG** | **0.15** | - | **2.65** | **5.04** |
| **Raccoon IgG** | **0.40** | + | **4.76** | **9.83** |
| **Ostrich IgY** | **0.21** | - | **3.28** | **4.50** |
| **Pig IgG** | **0.95** | +++ | **3.80** | **6.43** |
| **Mouse IgG** | **1.06** | +++ | **4.47** | **7.98** |
| **Bovine IgG** | **0.11** | - | **3.77** | **4.97** |
| **Rabbit IgG** | **0.41** | + | **4.43** | **6.10** |
| **Chicken IgY** | **0.16** | - | **3.10** | **4.26** |
| **Cat IgG** | **0.14** | - | **4.08** | **6.20** |
| **Rat IgG** | **0.09** | - | **2.65** | **4.25** |

### SpL and SpLA sandwich ELISA

This ELISA is used to study the interactions between protein-L (SpL) and protein-LA (SpLA) with different immunoglobulin preparations from mammalian and avian species. The 96 well microtitre plate is coated overnight at 4°C with 2 μg/μl per well of SpL in carbonate-bicarbonate buffer pH 9.6. Then plate is treated with bovine serum albumin solution and washed 4X with PBS-Tween. 50 μl of immunoglobulins (1 mg/ml) is added and incubated for 1h at room temperature (RT) and the microplate is rewashed 4X with PBS-Tween. Then 50 μl of peroxidase-labeled SpLA conjugate diluted 1:5000 in PBS-non-fat milk is added to each well and incubated for 1h at RT. The plate is washed 4X with PBS-Tween. 50 μl of 4 mg/ml o-phenylenediamine solution (OPD) is added and the plate is incubated 15 minutes at RT in the for the development of colour and read in a microplate reader at 492 nm. A cut-off point is calculated as the mean of the optical density of negative controls times two. The cut-off point is 0.26.

This immunoassay like the just above characterizes by fewer interactions between the bacterial immunoglobulin-binding proteins and IgGs. Table 15 depicts the SpL-SpLA sandwich ELISA, where SpL does not react with immunoglobulins from species of donkey, horse, mule, skunk, ostrich, bovine, cat, and rat. It suggests that SpL may not be suitable for certain assays, such as the study of the presence of antibodies against certain zoonotic pathogens in a greater number of animals in a population [10].

**Table 15.** SpL and SpLA sandwich ELISA.

| **Immunoglobulins** | **XOD results** | **Binding affinity** | **Coefficient of variation (CV) intra-assay (%)** | **CV inter-assay (%)** |
| --- | --- | --- | --- | --- |
| **Mule IgG** | **0.15** | **-** | **3.85** | **5.50** |
| **Donkey IgG** | **0.12** | **-** | **3.46** | **4.00** |
| **Horse IgG** | **0.17** | **-** | **2.25** | **3.73** |
| **Dog IgG** | **0.38** | **+** | **3.30** | **4.68** |
| **Skunk IgG** | **0.20** | **-** | **4.01** | **7.55** |
| **Coyote IgG** | **0.15** | **-** | **2.65** | **5.04** |
| **Raccoon IgG** | **0.40** | **+** | **4.76** | **9.83** |
| **Ostrich IgY** | **0.20** | **-** | **3.28** | **4.50** |
| **Pig IgG** | **0.95** | **+++** | **3.80** | **6.43** |
| **Mouse IgG** | **1.06** | **+++** | **4.47** | **7.98** |
| **Bovine IgG** | **0.11** | **-** | **3.77** | **4.97** |
| **Rabbit IgG** | **0.41** | **+** | **4.43** | **6.10** |
| **Chicken IgY** | **0.13** | **-** | **3.10** | **4.26** |
| **Cat IgG** | **0.14** | **-** | **2.33** | **3.78** |
| **Rat IgG** | **0.18** | **-** | **2.86** | **3.90** |

### SpA and PLAG sandwich ELISA

This ELISA is used to study the interactions between Staphylococcal protein-A (SpA) and protein-LAG (PLAG) with different immunoglobulin preparations from mammalian and avian species. The 96 well microtitre plate is coated overnight at 4°C with 2 μg/μl per well of SpA in carbonate-bicarbonate buffer pH 9.6. Then plate is treated with bovine serum albumin solution and washed 4X with PBS-Tween. 50 μl of immunoglobulins (1 mg/ml) is added and incubated for 1h at room temperature and the microplate is rewashed 4X with PBS-Tween. Then 50 μl of peroxidase-labeled PLAG conjugate diluted 1:5000 in PBS-non-fat milk is added to each well and incubated for 1h at RT. The plate is washed 4X with PBS-Tween. 50 μl of 4 mg/ml o-phenylenediamine solution (OPD) is added and the plate is incubated 15 minutes at RT in the dark. The reaction is stopped with 50 μl of 3M H2SO4 solution. The plate is visually assessed for the development of colour and read in a microplate reader at 492 nm. A cut-off point is calculated as the mean of the optical density of negative controls times two. The cut-off point is 0.32.

In this ELISA SpA and PLAG interact strongly with IgGs from cats, rabbits, mice, pigs, raccoons, coyotes, skunks, and dogs as shown in Table 16. They react moderately with IgG from bovines, horse, donkeys, and mules.

**Table 16.** SpA and PLAG sandwich ELISA.

| Immunoglobulins | XOD results | Binding affinity | Coefficient of variation (CV) intra-assay (%) | CV inter-assay (%) |
| --- | --- | --- | --- | --- |
| Mule IgG | 0.76 | ++ | 4.18 | 7.28 |
| Donkey IgG | 0.80 | ++ | 3.78 | 5.95 |
| Horse IgG | 0.78 | ++ | 3.06 | 5.70 |
| Dog IgG | 1.10 | +++ | 2.79 | 4.86 |
| Skunk IgG | 1.25 | +++ | 4.05 | 6.37 |
| Coyote IgG | 1.19 | +++ | 3.80 | 6.21 |
| Raccoon IgG | 1.30 | +++ | 3.68 | 5.07 |
| Ostrich IgY | 0.53 | + | 4.86 | 7.10 |
| Pig IgG | 1.37 | +++ | 2.96 | 4.97 |
| Mouse IgG | 1.39 | +++ | 4.00 | 7.67 |
| Bovine IgG | 0.78 | ++ | 5.29 | 8.42 |
| Rabbit IgG | 1.04 | +++ | 4.76 | 6.60 |
| Chicken IgY | 0.16 | - | 3.50 | 5.73 |
| Cat IgG | 1.35 | +++ | 3.79 | 6.01 |
| Rat IgG | 0.38 | + | 4.86 | 6.52 |

### SpG and PLAG sandwich ELISA

This ELISA is used to study the interactions between Streptococcal protein-G (SpG) and PLAG (PLAG) with different immunoglobulin preparations from mammalian and avian species. The 96 well microtitre plate is coated overnight at 4°C with 2 μg/μl per well of SpG in carbonate-bicarbonate buffer pH 9.6. Then plate is treated with bovine serum albumin solution and washed 4X with PBS-Tween. 50 μl of immunoglobulins (1 mg/ml) is added and incubated for 1h at room temperature and the microplate is rewashed 4X with PBS-Tween. Then 50 μl of peroxidase-labeled PLAG conjugate diluted 1:5000 in PBS-non-fat milk is added to each well and incubated for 1h at RT. The plate is washed 4X with PBS-Tween. 50 μl of 4 mg/ml o-phenylenediamine solution (OPD) is added and the plate is incubated 15 minutes at RT in the dark. The reaction is stopped with 50 μl of 3M H2SO4 solution. The plate is visually assessed for the development of colour and read in a microplate reader at 492 nm. A cut-off point is calculated as the mean of the optical density of negative controls times two. The cut-off point is 0.30.

This immunoassay resembles the previous one, in this test SpG and PLAG interact strongly with IgG from species of rabbit, bovine, mouse, pig, skunk, dog, horse, donkey, and mule as shown in Table 17.

**Table 17.** SpG and PLAG sandwich ELISA.

| **Immunoglobulins** | **XOD results** | **Binding affinity** | **Coefficient of variation (CV) intra-assay (%)** | **CV inter-assay (%)** |
| --- | --- | --- | --- | --- |
| **Mule IgG** | **0.95** | **+++** | **3.25** | **4.79** |
| **Donkey IgG** | **1.05** | **+++** | **3.76** | **5.15** |
| **Horse IgG** | **1.10** | **+++** | **4.09** | **6.22** |
| **Dog IgG** | **1.35** | **+++** | **3.35** | **5.14** |
| **Skunk IgG** | **1.30** | **+++** | **3.78** | **5.53** |
| **Coyote IgG** | **0.80** | **++** | **4.04** | **6.03** |
| **Raccoon IgG** | **0.43** | **+** | **3.25** | **5.18** |
| **Ostrich IgY** | **0.36** | **+** | **3.80** | **5.23** |
| **Pig IgG** | **1.30** | **+++** | **3.95** | **6.35** |
| **Mouse IgG** | **1.35** | **+++** | **3.33** | **4.80** |
| **Bovine IgG** | **1.05** | **+++** | **3.56** | **4.96** |
| **Rabbit IgG** | **1.13** | **+++** | **3.07** | **5.35** |
| **Chicken IgY** | **0.15** | **-** | **3.45** | **5.67** |
| **Cat IgG** | **0.40** | **+** | **4.26** | **5.07** |
| **Rat IgG** | **0.80** | **++** | **3.20** | **5.50** |

### SpL and PLAG sandwich ELISA

This ELISA is used to study the interactions between Peptostreptococcal protein-L (SpL) and protein LAG (PLAG) with different immunoglobulin preparations from mammalian and avian species. The 96 well microtitre plate is coated overnight at 4°C with 2 μg/μl per well of SpL in carbonate-bicarbonate buffer pH 9.6. Then plate is treated with bovine serum albumin solution and washed 4X with PBS-Tween. 50 μl of immunoglobulins (1 mg/ml) is added and incubated for 1h at room temperature and then, the microplate is rewashed 4X with PBS-Tween. Then 50 μl of peroxidase-labeled PLAG conjugate diluted 1:5000 in PBS-non-fat milk is added to each well and incubated for 1h at RT. The plate is washed 4X with PBS-Tween. 50 μl of 4 mg/ml o-phenylenediamine solution (OPD) is added and the plate is incubated 15 minutes at RT in the dark. The reaction is stopped with 50 μl of 3M H2SO4 solution. The plate is visually assessed for the development of colour and read in a microplate reader at 492 nm. A cut-off point is calculated as the mean of the optical density of negative controls times two. The cut-off point was 0.28.

Table 18 shows the binding affinity of an ELISA, where the bacterial receptor SpL binds to only immunoglobulin G from fewer species of animals. It binds weakly to IgGs from rabbits, raccoons, and dogs and very strongly to IgGs of mice and pigs. SpL does not bind to the rest of the immunoglobulin panel. It supports the fact that in an ELISA the reagent bound to the solid phase is essential in immunodetection.

**Table 18.** SpL and PLAG sandwich ELISA.

| Immunoglobulins | XOD results | Binding affinity | Coefficient of variation (CV) intra-assay (%) | CV inter-assay (%) |
| --- | --- | --- | --- | --- |
| Mule IgG | 0.17 | - | 3.85 | 5.50 |
| Donkey IgG | 0.15 | - | 3.46 | 4.00 |
| Horse IgG | 0.17 | - | 3.14 | 4.74 |
| Dog IgG | 0.40 | + | 2.67 | 3.65 |
| Skunk IgG | 0.14 | - | 3.01 | 5.54 |
| Coyote IgG | 0.18 | - | 3.65 | 4.52 |
| Raccoon IgG | 0.38 | + | 5.76 | 7.83 |
| Ostrich IgY | 0.14 | - | 2.55 | 3.40 |
| Pig IgG | 0.75 | +++ | 3.86 | 7.06 |
| Mouse IgG | 0.90 | +++ | 2.89 | 4.98 |
| Bovine IgG | 0.15 | - | 2.66 | 4.80 |
| Rabbit IgG | 0.36 | + | 3.05 | 4.10 |
| Chicken IgY | 0.14 | - | 3.10 | 6.35 |
| Cat IgG | 0.14 | - | 4.14 | 7.95 |
| Rat IgG | 0.12 | - | 3.70 | 5.18 |

### SpLA and PLAG sandwich ELISA

This ELISA is used to study the interactions between protein LA (SpLA) and protein-LAG (PLAG) with different immunoglobulin preparations from mammalian and avian species. The 96 well microtitre plate is coated overnight at 4°C with 2 μg/μl per well of SpLA in carbonate-bicarbonate buffer pH 9.6. Then plate is treated with bovine serum albumin solution and washed 4X with PBS-Tween. 50 μl of immunoglobulins (1 mg/ml) is added and incubated for 1h at room temperature and the microplate is rewashed 4X with PBS-Tween. Then 50 μl of peroxidase-labeled PLAG conjugate diluted 1:5000 in PBS-non-fat milk is added to each well and incubated for 1h at RT. The plate is washed 4X with PBS-Tween. 50 μl of 4 mg/ml o-phenylenediamine solution (OPD) is added and the plate is incubated 15 minutes at RT in the dark. The reaction is stopped with 50 μl of 3M H2SO4 solution. The plate is visually assessed for the development of colour and read in a microplate reader at 492 nm. A cut-off point is calculated as the mean of the optical density of negative controls times two. The cut-off point is 0.30.

Table 19 shows SpLA and PLAG binding strongly to the panel of Ig molecules. They react strongly with antibodies from species of skunk, raccoon, pig, rat, mouse, rabbit, and cat.

**Table 19.** SpLA and PLAG sandwich ELISA.

| **Immunoglobulins** | **XOD results** | **Binding affinity** | **Coefficient of variation (CV) intra-assay (%)** | **CV inter-assay (%)** |
| --- | --- | --- | --- | --- |
| **Mule IgG** | **0.39** | **+** | **3.41** | **5.25** |
| **Donkey IgG** | **0.44** | **+** | **3.22** | **4.61** |
| **Horse IgG** | **0.48** | **+** | **3.45** | **5.09** |
| **Dog IgG** | **0.75** | **++** | **4.27** | **5.80** |
| **Skunk IgG** | **0.95** | **+++** | **3.26** | **4.60** |
| **Coyote IgG** | **0.68** | **++** | **3.89** | **5.18** |
| **Raccoon IgG** | **1.10** | **+++** | **4.03** | **5.79** |
| **Ostrich IgY** | **0.33** | **+** | **3.68** | **5.79** |
| **Pig IgG** | **1.35** | **+++** | **4.00** | **5.24** |
| **Mouse IgG** | **1.40** | **+++** | **3.08** | **4.09** |
| **Bovine IgG** | **0.78** | **++** | **3.85** | **5.43** |
| **Rabbit IgG** | **1.05** | **+++** | **4.09** | **5.12** |
| **Chicken IgY** | **0.15** | **-** | **3.53** | **4.67** |
| **Cat IgG** | **1.15** | **+++** | **3.36** | **5.04** |
| **Rat IgG** | **0.94** | **+++** | **4.19** | **6.07** |

### SpLG and PLAG sandwich ELISA

This ELISA is used to study the interactions between protein-LG (SpLG) and protein-LAG (PLAG) with different immunoglobulin preparations from mammalian and avian species. The 96 well microtitre plate is coated overnight at 4°C with 2 μg/μl per well of SpLG in carbonate-bicarbonate buffer pH 9.6. Then plate is treated with bovine serum albumin solution and washed 4X with PBS-Tween. 50 μl of immunoglobulins (1 mg/ml) is added and incubated for 1h at room temperature and the microplate is rewashed 4X with PBS-Tween. Then 50 μl of peroxidase-labeled PLAG conjugate diluted 1:5000 in PBS-non-fat milk is added to each well and incubated for 1h at RT. The plate is washed 4X with PBS-Tween. 50 μl of 4 mg/ml o-phenylenediamine solution (OPD) is added and the plate is incubated 15 minutes at RT in the dark. The reaction is stopped with 50 μl of 3M H2SO4 solution. The plate is visually assessed for the development of colour and read in a microplate reader at 492 nm. A cut-off point is calculated as the mean of the optical density of negative controls times two. The cut-off point is 0.32.

SpLG and PLAG react strongly with some Ig molecules from much species including horse, dog, rat, pig, and mouse as shown in Table 20. However, cat IgG does not react with one the bacterial proteins in this assay (SpLG).

**Table 20.** SpLG and PLAG sandwich ELISA.

| **Immunoglobulins** | **XOD results** | **Binding affinity** | **Coefficient of variation (CV) intra-assay (%)** | **CV inter-assay (%)** |
| --- | --- | --- | --- | --- |
| **Mule IgG** | **0.79** | **++** | **4.13** | **6.88** |
| **Donkey IgG** | **0.95** | **+++** | **4.96** | **8.75** |
| **Horse IgG** | **1.05** | **+++** | **3.95** | **7.02** |
| **Dog IgG** | **1.16** | **+++** | **4.21** | **7.18** |
| **Skunk IgG** | **0.97** | **+++** | **4.50** | **8.11** |
| **Coyote IgG** | **0.70** | **++** | **5.02** | **9.18** |
| **Raccoon IgG** | **0.80** | **++** | **3.04** | **5.80** |
| **Ostrich IgY** | **0.36** | **+** | **4.43** | **6.57** |
| **Pig IgG** | **1.45** | **+++** | **3.75** | **6.90** |
| **Mouse IgG** | **1.30** | **+++** | **5.23** | **9.67** |
| **Bovine IgG** | **1.25** | **+++** | **5.16** | **10.05** |
| **Rabbit IgG** | **0.80** | **+++** | **4.24** | **6.42** |
| **Chicken IgY** | **0.16** | **-** | **3.98** | **5.43** |
| **Cat IgG** | **0.17** | **-** | **4.14** | **7.28** |
| **Rat IgG** | **1.45** | **+++** | **4.60** | **8.25** |

### SpAG and PLAG sandwich ELISA

This ELISA is used to study the interaction between protein-AG (SpAG) and protein-LAG (PLAG) with different immunoglobulin preparations from mammalian and avian species. The 96 well microtitre plate is coated overnight at 4°C with 2 μg/μl per well of SpAG in carbonate-bicarbonate buffer pH 9.6. Then plate is treated with bovine serum albumin solution and washed 4X with PBS-Tween. 50 μl of immunoglobulins (1 mg/ml) is added and incubated for 1h at room temperature and the microplate is rewashed 4X with PBS-Tween. Then 50 μl of peroxidase-labeled PLAG conjugate diluted 1:5000 in PBS-non-fat milk is added to each well and incubated for 1h at RT. The plate is washed 4X with PBS-Tween. 50 μl of 4 mg/ml o-phenylenediamine solution (OPD) is added and the plate is incubated 15 minutes at RT in the dark. The reaction is stopped with 50 μl of 3M H2SO4 solution. The plate is visually assessed for the development of colour and read in a microplate reader at 492 nm. A cut-off point is calculated as the mean of the optical density of negative controls times two. The cut-off point is 0.28.

Table 21 depicts that all immunoglobulins react effectively with SpAG-PLAG, except the ostrich IgY that binds weakly and the chicken IgY that does not react with any of the bacterial proteins and it is used as a negative control in this panel of 35 ELISAs.

**Table 21.**
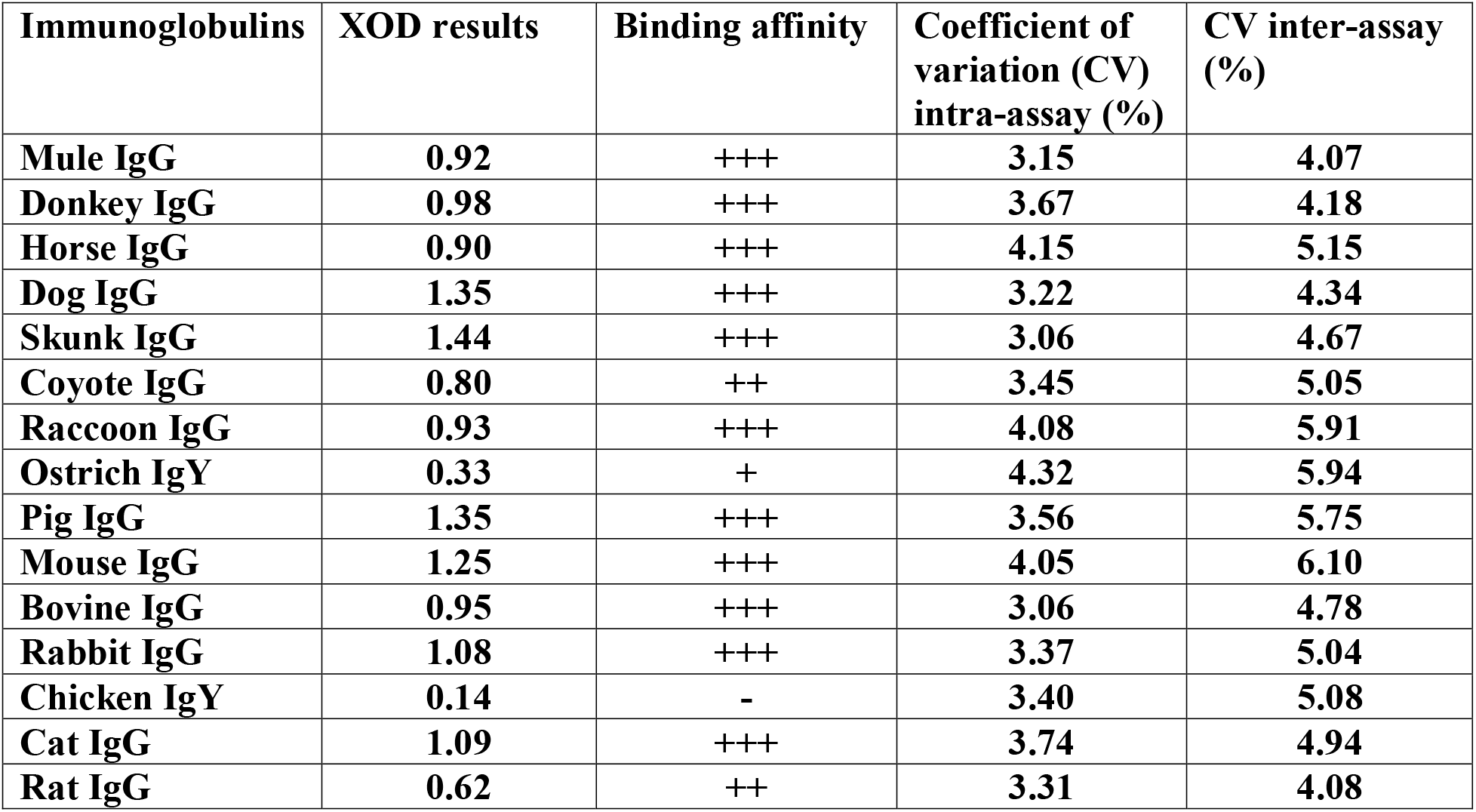
SpAG and PLAG sandwich ELISA.

| **Immunoglobulins** | **XOD results** | **Binding affinity** | **Coefficient of variation (CV) intra-assay (%)** | **CV inter-assay (%)** |
| --- | --- | --- | --- | --- |
| **Mule IgG** | **0.92** | **+++** | **3.15** | **4.07** |
| **Donkey IgG** | **0.98** | **+++** | **3.67** | **4.18** |
| **Horse IgG** | **0.90** | **+++** | **4.15** | **5.15** |
| **Dog IgG** | **1.35** | **+++** | **3.22** | **4.34** |
| **Skunk IgG** | **1.44** | **+++** | **3.06** | **4.67** |
| **Coyote IgG** | **0.80** | **++** | **3.45** | **5.05** |
| **Raccoon IgG** | **0.93** | **+++** | **4.08** | **5.91** |
| **Ostrich IgY** | **0.33** | **+** | **4.32** | **5.94** |
| **Pig IgG** | **1.35** | **+++** | **3.56** | **5.75** |
| **Mouse IgG** | **1.25** | **+++** | **4.05** | **6.10** |
| **Bovine IgG** | **0.95** | **+++** | **3.06** | **4.78** |
| **Rabbit IgG** | **1.08** | **+++** | **3.37** | **5.04** |
| **Chicken IgY** | **0.14** | **-** | **3.40** | **5.08** |
| **Cat IgG** | **1.09** | **+++** | **3.74** | **4.94** |
| **Rat IgG** | **0.62** | **++** | **3.31** | **4.08** |

### PLAG and PLAG sandwich ELISA

This ELISA is used to study the interactions of protein-LAG (PLAG) with different immunoglobulin preparations from mammalian and avian species. The 96 well microtitre plate is coated overnight at 4°C with 2 μg/μl per well of PLAG in carbonate-bicarbonate buffer pH 9.6. Then plate is treated with bovine serum albumin solution and washed 4X with PBS-Tween. 50 μl of immunoglobulins (1 mg/ml) is added and incubated for 1h at room temperature and the microplate is rewashed 4X with PBS-Tween. Then 50 μl of peroxidase-labeled PLAG conjugate diluted 1:5000 in PBS-non-fat milk is added to each well and incubated for 1h at RT. The plate is washed 4X with PBS-Tween. 50 μl of 4 mg/ml o-phenylenediamine solution (OPD) is added and the plate is incubated 15 minutes at RT in the dark. The reaction is stopped with 50 μl of 3M H2SO4 solution. The plate is visually assessed for the development of colour and read in a microplate reader at 492 nm. A cut-off point is calculated as the mean of the optical density of negative controls times two. The cut-off point is 0.32.

This is a very effective immunoassay. Fourteen out of 15 different immunoglobulins react with PLAG as shown in Table 22. A system like this brings the possibility or danger of causing steric hindrance. Nevertheless, it did not happen, and the fact that 86.6% of the immunoglobulins bound strongly to this hybrid protein proves it.

**Table 22.**
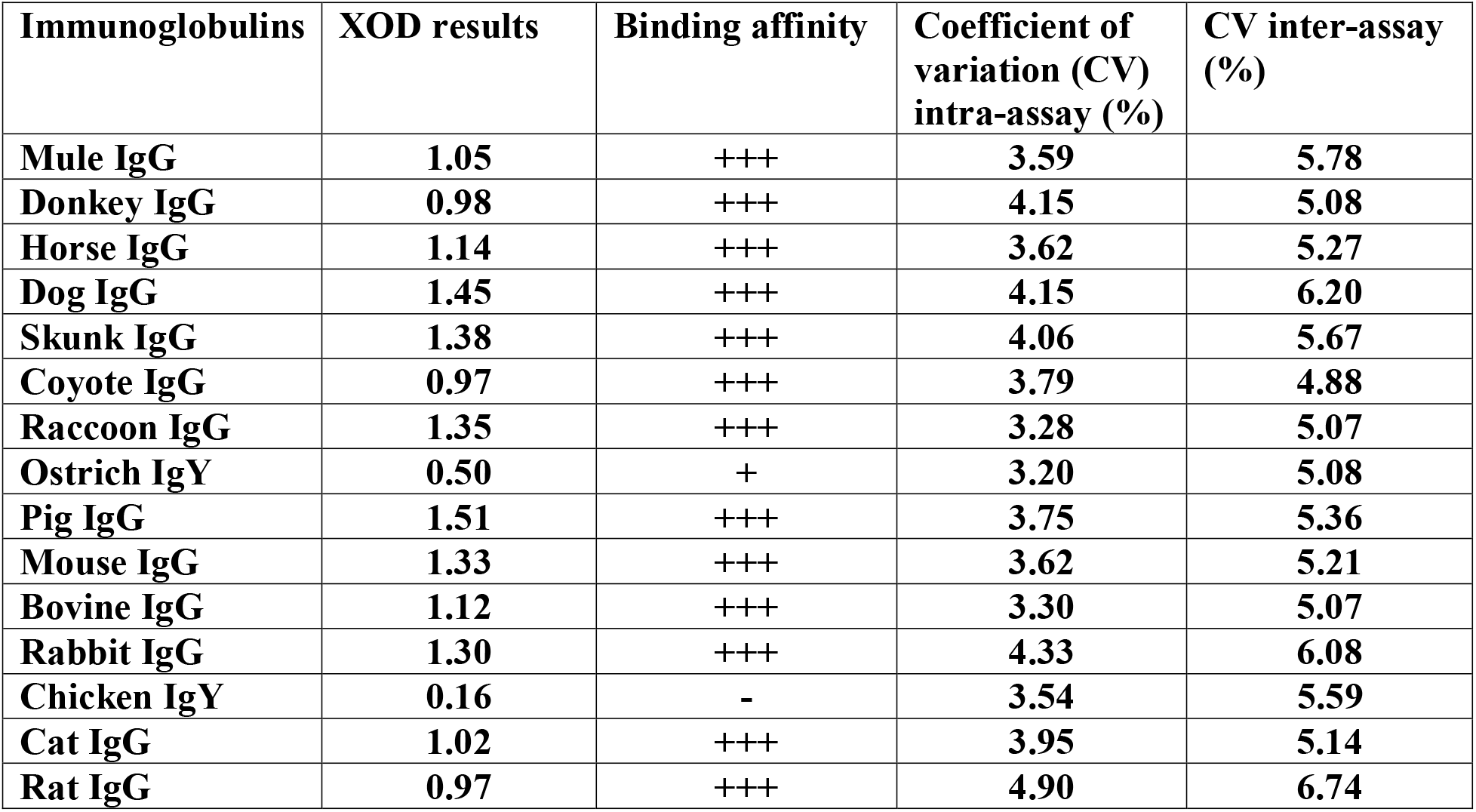
PLAG and PLAG sandwich ELISA.

### PLAG and SpL sandwich ELISA

This ELISA is used to study the interactions between protein-LAG (PLAG) and Peptostreptococcal protein-L (SpL) with different immunoglobulin preparations from mammalian and avian species. The 96 well microtitre plate is coated overnight at 4°C with 2 μg/μl per well of PLAG in carbonate-bicarbonate buffer pH 9.6. Then plate is treated with bovine serum albumin solution and washed 4X with PBS-Tween. 50 μl of immunoglobulins (1 mg/ml) is added and incubated for 1h at room temperature and then, the microplate is rewashed 4X with PBS-Tween. Then 50 μl of peroxidase-labeled SpL conjugate diluted 1:5000 in PBS-non-fat milk is added to each well and incubated for 1h at RT. The plate is washed 4X with PBS-Tween. 50 μl of 4 mg/ml o-phenylenediamine solution (OPD) is added and the plate is incubated 15 minutes at RT in the dark. The reaction is stopped with 50 μl of 3M H2SO4 solution. The plate is visually assessed for the development of colour and read in a microplate reader at 492 nm. A cut-off point is calculated as the mean of the optical density of negative controls times two. The cut-off point is 0.28.

Table 23 shows the PLAG-SpL sandwich ELISA is an interesting assay, because it is expected that since PLAG is involved, more higher affinities can be picked up. Unfortunately, this assay’s sensitivity is low because it uses a peroxidase conjugated SpL that fails to interact with many immunoglobulins.

**Table 23.** PLAG and SpL sandwich ELISA.

| **Immunoglobulins** | **XOD results** | **Binding affinity** | **Coefficient of variation (CV) intra-assay (%)** | **CV inter-assay (%)** |
| --- | --- | --- | --- | --- |
| **Mule IgG** | **0.23** | - | **3.85** | **5.50** |
| **Donkey IgG** | **0.15** | - | **3.46** | **4.02** |
| **Horse IgG** | **0.17** | - | **3.14** | **4.74** |
| **Dog IgG** | **0.40** | + | **2.67** | **3.65** |
| **Skunk IgG** | **0.14** | - | **3.01** | **5.54** |
| **Coyote IgG** | **0.18** | - | **3.65** | **4.54** |
| **Raccoon IgG** | **0.38** | + | **2.76** | **4.83** |
| **Ostrich IgY** | **0.14** | - | **3.35** | **6.40** |
| **Pig IgG** | **0.75** | +++ | **2.86** | **4.82** |
| **Mouse IgG** | **0.90** | +++ | **2.80** | **3.77** |
| **Bovine IgG** | **0.20** | - | **2.88** | **3.50** |
| **Rabbit IgG** | **0.39** | + | **3.05** | **5.10** |
| **Chicken IgY** | **0.14** | - | **2.95** | **3.89** |
| **Cat IgG** | **0.12** | - | **3.08** | **5.70** |
| **Rat IgG** | **0.12** | - | **2.88** | **4.60** |

### PLAG and SpA sandwich ELISA

This ELISA is used to study the interactions between protein-LAG (PLAG) and Staphylococcal protein-A (SpA) with different immunoglobulin preparations from mammalian and avian species. The 96 well microtitre plate is coated overnight at 4°C with 2 μg/μl per well of PLAG in carbonate-bicarbonate buffer pH 9.6. Then plate is treated with bovine serum albumin solution and washed 4X with PBS-Tween. 50 μl of immunoglobulins (1 mg/ml) is added and incubated for 1h at room temperature and the microplate is rewashed 4X with PBS-Tween. Then 50 μl of peroxidase-labeled SpA conjugate diluted 1:5000 in PBS-non-fat milk is added to each well and incubated for 1h at RT. The plate is washed 4X with PBS-Tween. 50 μl of 4 mg/ml o-phenylenediamine solution (OPD) is added and the plate is incubated 15 minutes at RT in the dark. The reaction is stopped with 50 μl of 3M H2SO4 solution. The plate is visually assessed for the development of colour and read in a microplate reader at 492 nm. A cut-off point is calculated as the mean of the optical density of negative controls times two. The cut-off point is 0.30.

Table 24 shows an ELISA where SpA interacts strongly with the 46.6% of the panel of immunoglobulins (7 out of 15). SpA binds moderately to IgG of species of mule, donkey, coyote, and horse.

**Table 24.** PLAG and SpA sandwich ELISA.

| **Immunoglobulins** | **XOD results** | **Binding affinity** | **Coefficient of variation (CV) intra-assay (%)** | **CV inter-assay (%)** |
| --- | --- | --- | --- | --- |
| **Mule IgG** | **0.65** | **++** | **4.22** | **8.15** |
| **Donkey IgG** | **0.70** | **++** | **4.57** | **7.90** |
| **Horse IgG** | **0.74** | **++** | **4.05** | **6.70** |
| **Dog IgG** | **1.20** | **+++** | **3.67** | **6.86** |
| **Skunk IgG** | **1.27** | **+++** | **4.44** | **8.19** |
| **Coyote IgG** | **0.80** | **++** | **3.56** | **5.80** |
| **Raccoon IgG** | **1.25** | **+++** | **5.01** | **9.33** |
| **Ostrich IgY** | **0.45** | **+** | **3.79** | **7.10** |
| **Pig IgG** | **1.30** | **+++** | **4.57** | **7.97** |
| **Mouse IgG** | **1.42** | **+++** | **3.06** | **5.75** |
| **Bovine IgG** | **0.45** | **+** | **4.90** | **8.74** |
| **Rabbit IgG** | **1.13** | **+++** | **3.76** | **6.03** |
| **Chicken IgY** | **0.15** | **-** | **3.55** | **5.91** |
| **Cat IgG** | **1.17** | **+++** | **3.69** | **6.54** |
| **Rat IgG** | **0.35** | **+** | **4.68** | **7.05** |

### PLAG and SpG sandwich ELISA

This ELISA is used to study the interactions between protein-LAG (PLAG) and Streptococcal protein-G (SpG) with different immunoglobulin preparations from mammalian and avian species. The 96 well microtitre plate is coated overnight at 4°C with 2 μg/μl per well of PLAG in carbonate-bicarbonate buffer pH 9.6. Then plate is treated with bovine serum albumin solution and washed 4X with PBS-Tween. 50 μl of immunoglobulins (1 mg/ml) is added and incubated for 1h at room temperature and the microplate is rewashed 4X with PBS-Tween. Then 50 μl of peroxidase-labeled SpG conjugate diluted 1:5000 in PBS-non-fat milk is added to each well and incubated for 1h at RT. The plate is washed 4X with PBS-Tween. 50 μl of 4 mg/ml o-phenylenediamine solution (OPD) is added and the plate is incubated 15 minutes at RT in the dark. The reaction is stopped with 50 μl of 3M H2SO4 solution. The plate is visually assessed for the development of colour and read in a microplate reader at 492 nm. A cut-off point is calculated as the mean of the optical density of negative controls times two. The cut-off point is 0.30.

Table 25 shows a very powerful assay where SpG interact strongly with 66.6% of the immunoglobulin panel (10 out of 15). SpG does not react with chicken IgY and cat IgG.

**Table 25.** PLAG and SpG sandwich ELISA.

| **Immunoglobulins** | **XOD results** | **Binding affinity** | **Coefficient of variation (CV) intra-assay (%)** | **CV inter-assay (%)** |
| --- | --- | --- | --- | --- |
| **Mule IgG** | **1.03** | **+++** | **2.83** | **5.22** |
| **Donkey IgG** | **1.10** | **+++** | **2.95** | **4.89** |
| **Horse IgG** | **1.15** | **+++** | **2.86** | **5.65** |
| **Dog IgG** | **1.50** | **+++** | **3.05** | **6.38** |
| **Skunk IgG** | **1.44** | **+++** | **3.67** | **6.15** |
| **Coyote IgG** | **0.75** | **++** | **4.80** | **6.76** |
| **Raccoon IgG** | **0.40** | **+** | **2.89** | **5.73** |
| **Ostrich IgY** | **0.36** | **+** | **3.49** | **6.05** |
| **Pig IgG** | **1.53** | **+++** | **3.37** | **5.88** |
| **Mouse IgG** | **1.48** | **+++** | **2.81** | **4.80** |
| **Bovine IgG** | **0.99** | **+++** | **3.41** | **5.75** |
| **Rabbit IgG** | **1.02** | **+++** | **4.05** | **7.07** |
| **Chicken IgY** | **0.15** | **-** | **2.81** | **4.73** |
| **Cat IgG** | **0.15** | **-** | **3.08** | **5.14** |
| **Rat IgG** | **1.28** | **+++** | **3.90** | **6.01** |

### PLAG and SpLG sandwich ELISA

This ELISA is used to study the interactions between protein-LAG (PLAG) and protein-LG (SpLG) with different immunoglobulin preparations from mammalian species. The 96 well microtitre plate is coated overnight at 4°C with 2 μg/μl per well of PLAG in carbonate-bicarbonate buffer pH 9.6. Then plate is treated with bovine serum albumin solution and washed 4X with PBS-Tween. 50 μl of immunoglobulins (1 mg/ml) is added and incubated for 1h at room temperature and the microplate is rewashed 4X with PBS-Tween. Then 50 μl of peroxidase-labeled SpLG conjugate diluted 1:5000 in PBS-non-fat milk is added to each well and incubated for 1h at RT. The plate is washed 4X with PBS-Tween. 50 μl of 4 mg/ml o-phenylenediamine solution (OPD) is added and the plate is incubated 15 minutes at RT in the dark. The reaction is stopped with 50 μl of 3M H2SO4 solution. The plate is visually assessed for the development of colour and read in a microplate reader at 492 nm. A cut-off point is calculated as the mean of the optical density of negative controls times two. The cut-off point is 0.34.

Table 26 depicts the PLAG and SpLG sandwich ELISA. It is an interesting assay because in its solid phase it binds to 14 out of 15 immunoglobulins. On the other hand, the conjugate (SpLG-HRP) does not interact with the two avian immunoglobulins, neither the cat IgG, nor very weakly react with the coyote IgG. However, this system interacts strongly with IgG from species of horse, pig, rat, among other species.

**Table 26.** PLAG and SpLG sandwich ELISA.

| **Immunoglobulins** | **XOD results** | **Binding affinity** | **Coefficient of variation (CV) intra-assay (%)** | **CV inter-assay (%)** |
| --- | --- | --- | --- | --- |
| **Mule IgG** | **1.10** | **+++** | **3.55** | **7.04** |
| **Donkey IgG** | **1.22** | **+++** | **3.65** | **6.19** |
| **Horse IgG** | **1.28** | **+++** | **4.24** | **6.86** |
| **Dog IgG** | **1.30** | **+++** | **3.15** | **5.75** |
| **Skunk IgG** | **1.35** | **+++** | **4.28** | **6.30** |
| **Coyote IgG** | **0.48** | **+** | **3.70** | **6.05** |
| **Raccoon IgG** | **0.70** | **++** | **4.08** | **8.45** |
| **Ostrich IgY** | **0.20** | **-** | **3.66** | **6.71** |
| **Pig IgG** | **1.35** | **+++** | **3.58** | **5.99** |
| **Mouse IgG** | **1.45** | **+++** | **3.69** | **6.43** |
| **Bovine IgG** | **1.08** | **+++** | **3.54** | **5.07** |
| **Rabbit IgG** | **1.14** | **+++** | **3.78** | **6.09** |
| **Chicken IgY** | **0.17** | **-** | **3.34** | **5.70** |
| **Cat IgG** | **0.11** | **-** | **3.46** | **6.28** |
| **Rat IgG** | **1.25** | **+++** | **3.57** | **7.07** |

### PLAG and SpLA sandwich ELISA

This ELISA is used to study the interactions between protein-LAG (PLAG) and protein-LA (SpLA) with different immunoglobulin preparations from mammalian and avian species. The 96 well microtitre plate is coated overnight at 4°C with 2 μg/μl per well of PLAG in carbonate-bicarbonate buffer pH 9.6. Then plate is treated with bovine serum albumin solution and washed 4X with PBS-Tween. 50 μl of immunoglobulins (1 mg/ml) is added and incubated for 1h at room temperature and the microplate is rewashed 4X with PBS-Tween. Then 50 μl of peroxidase-labeled SpLA conjugate diluted 1:5000 in PBS-non-fat milk is added to each well and incubated for 1h at RT. The plate is washed 4X with PBS-Tween. 50 μl of 4 mg/ml o-phenylenediamine solution (OPD) is added and the plate is incubated 15 minutes at RT in the dark. The reaction is stopped with 50 μl of 3M H2SO4 solution. The plate is visually assessed for the development of colour and read in a microplate reader at 492 nm. A cut-off point is calculated as the mean of the optical density of negative controls times two. The cut-off point is 0.32.

Table 27 shows the binding affinity of SpLA and PLAG to the immunoglobulin panel. They interact effectively with IgGs of cats, rabbits, mice, pigs, raccoons, skunks, and dogs.

**Table 27.** PLAG and SpLA sandwich ELISA.

| Immunoglobulins | XOD results | Binding affinity | Coefficient of variation (CV) intra-assay (%) | CV inter-assay (%) |
| --- | --- | --- | --- | --- |
| **Mule IgG** | **0.68** | ++ | **3.55** | **6.72** |
| **Donkey IgG** | **0.75** | ++ | **4.88** | **7.77** |
| **Horse IgG** | **0.75** | ++ | **3.85** | **5.06** |
| **Dog IgG** | **1.15** | +++ | **4.08** | **8.19** |
| **Skunk IgG** | **1.00** | +++ | **3.45** | **6.13** |
| **Coyote IgG** | **0.80** | ++ | **4.60** | **7.36** |
| **Raccoon IgG** | **1.15** | +++ | **3.85** | **5.09** |
| **Ostrich IgY** | **0.42** | + | **3.75** | **5.50** |
| **Pig IgG** | **1.26** | +++ | **4.65** | **6.22** |
| **Mouse IgG** | **1.31** | +++ | **2.96** | **5.18** |
| **Bovine IgG** | **0.80** | ++ | **3.19** | **5.58** |
| **Rabbit IgG** | **0.98** | +++ | **4.34** | **7.20** |
| **Chicken IgY** | **0.16** | - | **3.08** | **5.35** |
| **Cat IgG** | **1.25** | +++ | **5.24** | **8.64** |
| **Rat IgG** | **0.35** | + | **4.50** | **7.13** |

### Table 28. PLAG and SpAG sandwich ELISA

This ELISA is used to study the interactions between protein-LAG (PLAG) and protein-AG (SpAG) with different immunoglobulin preparations from mammalian and avian species. The 96 well microtitre plate is coated overnight at 4°C with 2 μg/μl per well of PLAG in carbonate-bicarbonate buffer pH 9.6. Then plate is treated with bovine serum albumin solution and washed 4X with PBS-Tween. 50 μl of immunoglobulins (1 mg/ml) is added and incubated for 1h at room temperature and the microplate is rewashed 4X with PBS-Tween. Then 50 μl of peroxidase-labeled SpAG conjugate diluted 1:5000 in PBS-non-fat milk is added to each well and incubated for 1h at RT. The plate is washed 4X with PBS-Tween. 50 μl of 4 mg/ml o-phenylenediamine solution (OPD) is added and the plate is incubated 15 minutes at RT in the dark. The reaction is stopped with 50 μl of 3M H2SO4 solution. The plate is visually assessed for the development of colour and read in a microplate reader at 492 nm. A cut-off point is calculated as the mean of the optical density of negative controls times two. The cut-off point is 0.36.

Table 28 represents an effective immunoassay capable of strongly interacting with IgGs of many species. Twelve out of 15 immunoglobulins (80%) bind strongly to SpAG.

**Table 28.** PLAG and SpAG sandwich ELISA.

| **Immunoglobulins** | **XOD results** | **Binding affinity** | **Coefficient of variation (CV) intra-assay (%)** | **CV inter-assay (%)** |
| --- | --- | --- | --- | --- |
| **Mule IgG** | **1.25** | **+++** | **3.67** | **6.63** |
| **Donkey IgG** | **1.28** | **+++** | **2.53** | **5.29** |
| **Horse IgG** | **1.35** | **+++** | **4.45** | **8.15** |
| **Dog IgG** | **1.50** | **+++** | **3.20** | **5.02** |
| **Skunk IgG** | **1.44** | **+++** | **3.09** | **5.79** |
| **Coyote IgG** | **1.11** | **+++** | **4.88** | **7.66** |
| **Raccoon IgG** | **0.80** | **++** | **3.28** | **5.55** |
| **Ostrich IgY** | **0.37** | **+** | **3.13** | **6.09** |
| **Pig IgG** | **1.55** | **+++** | **5.24** | **8.46** |
| **Mouse IgG** | **1.60** | **+++** | **3.29** | **4.97** |
| **Bovine IgG** | **1.07** | **+++** | **3.18** | **5.06** |
| **Rabbit IgG** | **1.16** | **+++** | **3.77** | **6.01** |
| **Chicken IgY** | **0.18** | **-** | **2.85** | **4.79** |
| **Cat IgG** | **0.95** | **+++** | **4.45** | **7.13** |
| **Rat IgG** | **1.05** | **+++** | **3.43** | **7.05** |

### SpA Direct ELISA

This ELISA is used to study the interactions of Staphylococcal protein-A (SpA) with diverse immunoglobulins. The 96 well microtitre plate is coated overnight at 4°C with 1 μg/μl per well of purified immunoglobulins in carbonate-bicarbonate buffer pH 9.6. Then, plate is treated with bovine serum albumin solution and washed 4X with PBS-Tween. Then, 50 μl of peroxidase-labeled-protein-A conjugate diluted 1:3000 in PBS-non-fat milk is added to each well and incubated for 1.30h at RT. After that, the plate is washed 4X with PBS-Tween. Pipette 50 μl of 3,3’,5,5’ - tetramethylbenzidine (TMB; Sigma-Aldrich) to each well. The reaction is stopped with 50 μl of 3M H2SO4 solution. The plate is visually assessed for the development of colour and read in a microplate reader at 450 nm. A cut-off point is calculated as the mean of the optical density of negative controls times two. The cut-off point was 0.28.

Table 29 shows SpA that binds strongly to the 46.66% of the panel of immunoglobulins. However, it binds moderately to IgGs of horses, coyotes, and bovines.

**Table 29.** SpA Direct ELISA.

| **Immunoglobulins** | **XOD results** | **Binding affinity** | **Coefficient of variation (CV) intra-assay (%)** | **CV inter-assay (%)** |
| --- | --- | --- | --- | --- |
| **Mule IgG** | **0.37** | **+** | **3.03** | **5.20** |
| **Donkey IgG** | **0.43** | **+** | **3.75** | **7.35** |
| **Horse IgG** | **0.52** | **++** | **4.32** | **6.08** |
| **Dog IgG** | **0.95** | **+++** | **3.22** | **5.07** |
| **Skunk IgG** | **0.90** | **+++** | **5.05** | **8.37** |
| **Coyote IgG** | **0.61** | **++** | **3.89** | **6.04** |
| **Raccoon IgG** | **0.96** | **+++** | **4.48** | **7.12** |
| **Ostrich IgY** | **0.35** | **+** | **3.75** | **6.18** |
| **Pig IgG** | **0.95** | **+++** | **3.87** | **5.75** |
| **Mouse IgG** | **0.98** | **+++** | **2.95** | **4.18** |
| **Bovine IgG** | **0.60** | **++** | **3.26** | **5.05** |
| **Rabbit IgG** | **0.90** | **+++** | **3.74** | **6.35** |
| **Chicken IgY** | **0.14** | **-** | **2.58** | **5.19** |
| **Cat IgG** | **1.05** | **+++** | **5.16** | **9.13** |
| **Rat IgG** | **0.39** | **+** | **3.80** | **6.15** |

### SpG Direct ELISA

This ELISA is used to study the interactions of Streptococcal protein-G (SpG) with diverse immunoglobulins. The 96 well microtitre plate is coated overnight at 4°C with 1 μg/μl per well of purified immunoglobulins in carbonate-bicarbonate buffer pH 9.6. Then, plate is treated with bovine serum albumin solution and washed 4X with PBS-Tween. Then, 50 μl of peroxidase-labeled-protein-G conjugate diluted 1:3000 in PBS-non-fat milk is added to each well and incubated for 1.30h at RT. After that, the plate is washed 4X with PBS-Tween. Pipette 50 μl of 3,3’,5,5’ - tetramethylbenzidine (TMB; Sigma-Aldrich) to each well. The reaction is stopped with 50 μl of 3M H2SO4 solution. The plate is visually assessed for the development of colour and read in a microplate reader at 450 nm. A cut-off point is calculated as the mean of the optical density of negative controls times two. The cut-off point is 0.28.

Table 30 shows a direct ELISA where SpG binds strongly to 53.33% of the immunoglobulin panel (8 out of 15 Igs).

**Table 30.** SpG Direct ELISA.

| **Immunoglobulins** | **XOD results** | **Binding affinity** | **Coefficient of variation (CV) intra-assay (%)** | **CV inter-assay (%)** |
| --- | --- | --- | --- | --- |
| **Mule IgG** | **0.70** | **++** | **3.86** | **6.13** |
| **Donkey IgG** | **0.75** | **++** | **3.50** | **5.84** |
| **Horse IgG** | **0.87** | **+++** | **4.12** | **7.06** |
| **Dog IgG** | **0.95** | **+++** | **3.59** | **6.61** |
| **Skunk IgG** | **0.90** | **+++** | **4.03** | **7.35** |
| **Coyote IgG** | **0.35** | **+** | **4.29** | **8.08** |
| **Raccoon IgG** | **0.33** | **+** | **3.67** | **5.06** |
| **Ostrich IgY** | **0.07** | **-** | **5.25** | **8.66** |
| **Pig IgG** | **1.11** | **+++** | **3.44** | **6.15** |
| **Mouse IgG** | **1.05** | **+++** | **4.04** | **6.57** |
| **Bovine IgG** | **0.95** | **+++** | **3.94** | **7.32** |
| **Rabbit IgG** | **0.88** | **+++** | **4.68** | **8.02** |
| **Chicken IgY** | **0.07** | **-** | **2.95** | **4.74** |
| **Cat IgG** | **0.14** | **-** | **3.52** | **5.08** |
| **Rat IgG** | **0.96** | **+++** | **3.67** | **5.86** |

### SpL Direct ELISA

This ELISA is used to study the interactions of Peptostreptococcal protein-L (SpL) with diverse immunoglobulins. The 96 well microtitre plate is coated overnight at 4°C with 1 μg/μl per well of purified immunoglobulins in carbonate-bicarbonate buffer pH 9.6. Then, plate is treated with bovine serum albumin solution and washed 4X with PBS-Tween. Then, 50 μl of peroxidase-labeled-protein-L conjugate diluted 1:3000 in PBS-non-fat milk is added to each well and incubated for 1.30h at RT. After that, the plate is washed 4X with PBS-Tween. Pipette 50 μl of 3,3’,5,5’ - tetramethylbenzidine (TMB; Sigma-Aldrich) to each well. The reaction is stopped with 50 μl of 3M H2SO4 solution. The plate is visually assessed for the development of colour and read in a microplate reader at 450 nm. A cut-off point is calculated as the mean of the optical density of negative controls times two. The cut-off point is 0.24.

Table 31 shows the direct ELISA where SpL strongly bind to 13.33% of the mammalian immunoglobulins. It does not bind to 66.6% of the panel of immunoglobulins.

**Table 31.** SpL Direct ELISA.

| **Immunoglobulins** | **XOD results** | **Binding affinity** | **Coefficient of variation (CV) intra-assay (%)** | **CV inter-assay (%)** |
| --- | --- | --- | --- | --- |
| **Mule IgG** | **0.09** | **-** | **2.84** | **4.41** |
| **Donkey IgG** | **0.10** | **-** | **2.50** | **4.15** |
| **Horse IgG** | **0.15** | **-** | **2.78** | **3.85** |
| **Dog IgG** | **0.33** | **+** | **2.36** | **3.60** |
| **Skunk IgG** | **0.14** | **-** | **3.01** | **5.73** |
| **Coyote IgG** | **0.14** | **-** | **2.80** | **4.08** |
| **Raccoon IgG** | **0.43** | **+** | **3.99** | **6.06** |
| **Ostrich IgY** | **0.09** | **-** | **2.54** | **3.07** |
| **Pig IgG** | **0.90** | **+++** | **3.80** | **7.43** |
| **Mouse IgG** | **0.98** | **+++** | **5.47** | **8.98** |
| **Bovine IgG** | **0.08** | **-** | **3.58** | **5.20** |
| **Rabbit IgG** | **0.34** | **+** | **4.67** | **7.23** |
| **Chicken IgY** | **0.06** | **-** | **2.40** | **4.65** |
| **Cat IgG** | **0.12** | **-** | **4.03** | **8.79** |
| **Rat IgG** | **0.08** | **-** | **3.15** | **5.02** |

### SpLA Direct ELISA

This ELISA is used to study the interactions of protein-LA (SpLA) with diverse immunoglobulins. The 96 well microtitre plate is coated overnight at 4°C with 1 μg/μl per well of purified immunoglobulins in carbonate-bicarbonate buffer pH 9.6. Then, plate is treated with bovine serum albumin solution and washed 4X with PBS-Tween. Then, 50 μl of peroxidase-labeled-protein-LA conjugate diluted 1:3000 in PBS-non-fat milk is added to each well and incubated for 1.30h at RT. After that, the plate is washed 4X with PBS-Tween. Pipette 50 μl of 3,3’,5,5’ - tetramethylbenzidine (TMB; Sigma-Aldrich) to each well. The reaction is stopped with 50 μl of 3M H2SO4 solution. The plate is visually assessed for the development of colour and read in a microplate reader at 450 nm. A cut-off point is calculated as the mean of the optical density of negative controls times two. The cut-off point is 0.30.

It depicts the direct SpLA ELISA, where the bacterial protein binds strongly to immunoglobulins of various species of animals including dog, raccoon, and rabbit. As shown in table 32, indeed, it does not bind to the negative control: the chicken IgY. SpLA is a very versatile reagent that strongly binds to immunoglobulins from many animal species [29].

**Table 32.** SpLA Direct ELISA.

| **Immunoglobulins** | **XOD results** | **Binding affinity** | **Coefficient of variation (CV) intra-assay (%)** | **CV inter-assay (%)** |
| --- | --- | --- | --- | --- |
| **Mule IgG** | **0.34** | **+** | **3.53** | **5.05** |
| **Donkey IgG** | **0.41** | **+** | **3.88** | **5.61** |
| **Horse IgG** | **0.68** | **++** | **3.67** | **5.08** |
| **Dog IgG** | **0.95** | **+++** | **4.58** | **6.37** |
| **Skunk IgG** | **0.91** | **+++** | **3.85** | **5.03** |
| **Coyote IgG** | **0.63** | **++** | **4.77** | **7.18** |
| **Raccoon IgG** | **0.98** | **+++** | **3.75** | **5.89** |
| **Ostrich IgY** | **0.38** | **+** | **4.54** | **7.17** |
| **Pig IgG** | **1.06** | **+++** | **5.06** | **8.56** |
| **Mouse IgG** | **1.15** | **+++** | **3.05** | **6.40** |
| **Bovine IgG** | **0.46** | **+** | **4.64** | **5.98** |
| **Rabbit IgG** | **0.96** | **+++** | **3.87** | **5.82** |
| **Chicken IgY** | **0.15** | **-** | **2.35** | **4.80** |
| **Cat IgG** | **0.85** | **+++** | **3.36** | **5.70** |
| **Rat IgG** | **0.35** | **+** | **4.11** | **6.16** |

### SpLG Direct ELISA

This ELISA is used to study the interactions of protein-LG (SpLG) with diverse immunoglobulins. The 96 well microtitre plate is coated overnight at 4°C with 1 μg/μl per well of purified immunoglobulins in carbonate-bicarbonate buffer pH 9.6. Then, plate is treated with bovine serum albumin solution and washed 4X with PBS-Tween. Then, 50 .μl of peroxidase-labeled-protein-LG conjugate diluted 1:3000 in PBS-non-fat milk is added to each well and incubated for 1.30h at RT. After that, the plate is washed 4X with PBS-Tween. Pipette 50 μl of 3,3’,5,5’ - tetramethylbenzidine (TMB; Sigma-Aldrich) to each well. The reaction is stopped with 50 μl of 3M H2SO4 solution. The plate is visually assessed for the development of colour and read in a microplate reader at 450 nm. A cut-off point is calculated as the mean of the optical density of negative controls times two. The cut-off point is 0.30.

Table 33 depicts the indirect ELISA, where SpLG strongly interacts with 46.66% of the panel of immunoglobulins. It has been shown to be a versatile IgG-binding reagent. It bound to mouse and rat IgG, and many other immunoglobulins and antibody fragments [30].

**Table 33.**
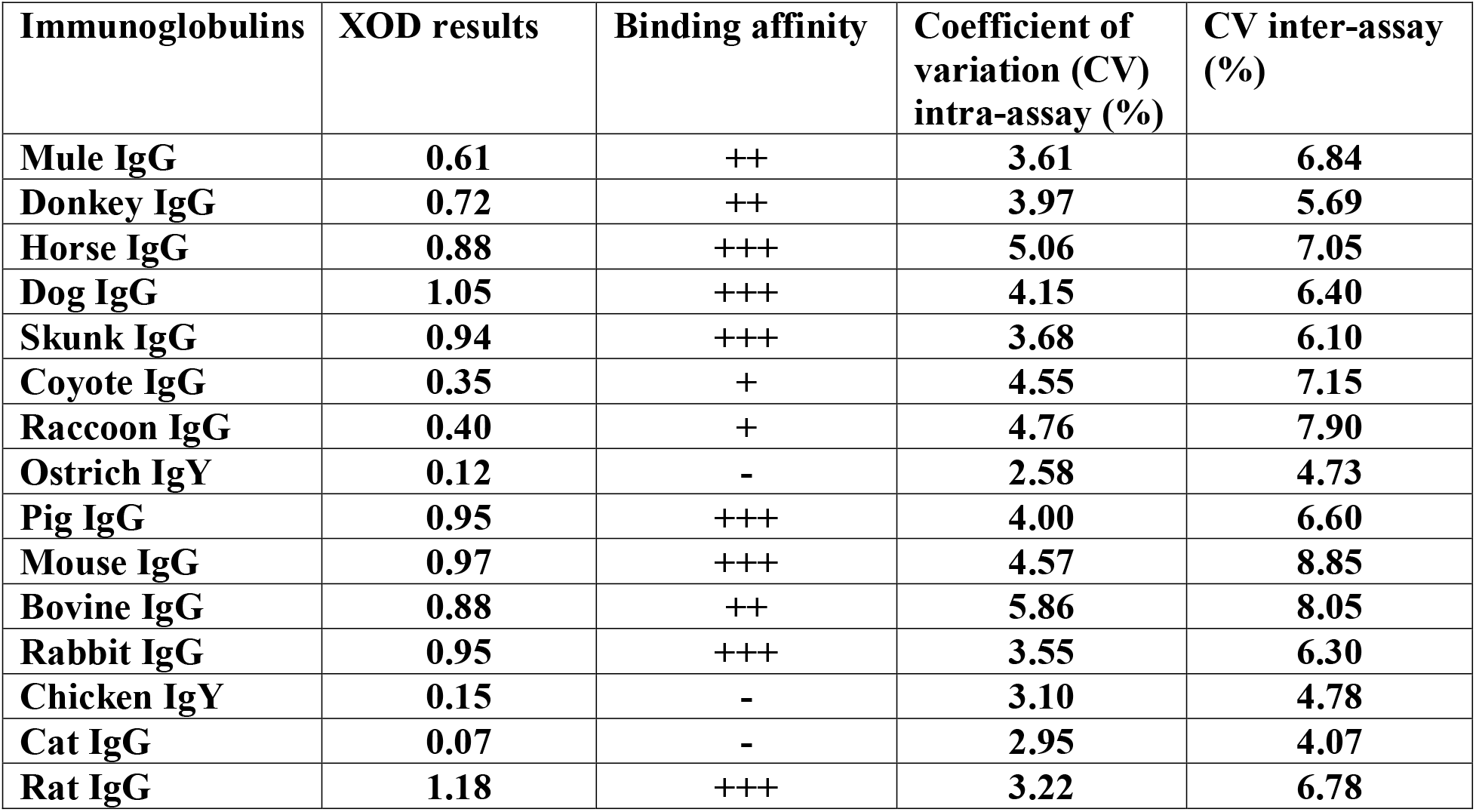
SpLG Direct ELISA.

| **Immunoglobulins** | **XOD results** | **Binding affinity** | **Coefficient of variation (CV) intra-assay (%)** | **CV inter-assay (%)** |
| --- | --- | --- | --- | --- |
| **Mule IgG** | **0.61** | **++** | **3.61** | **6.84** |
| **Donkey IgG** | **0.72** | **++** | **3.97** | **5.69** |
| **Horse IgG** | **0.88** | **+++** | **5.06** | **7.05** |
| **Dog IgG** | **1.05** | **+++** | **4.15** | **6.40** |
| **Skunk IgG** | **0.94** | **+++** | **3.68** | **6.10** |
| **Coyote IgG** | **0.35** | **+** | **4.55** | **7.15** |
| **Raccoon IgG** | **0.40** | **+** | **4.76** | **7.90** |
| **Ostrich IgY** | **0.12** | **-** | **2.58** | **4.73** |
| **Pig IgG** | **0.95** | **+++** | **4.00** | **6.60** |
| **Mouse IgG** | **0.97** | **+++** | **4.57** | **8.85** |
| **Bovine IgG** | **0.88** | **++** | **5.86** | **8.05** |
| **Rabbit IgG** | **0.95** | **+++** | **3.55** | **6.30** |
| **Chicken IgY** | **0.15** | **-** | **3.10** | **4.78** |
| **Cat IgG** | **0.07** | **-** | **2.95** | **4.07** |
| **Rat IgG** | **1.18** | **+++** | **3.22** | **6.78** |

### SpAG Direct ELISA

This ELISA is used to study the interactions of protein-AG (SpAG) with diverse immunoglobulins. The 96 well microtitre plate is coated overnight at 4°C with 1 μg/μl per well of purified immunoglobulins in carbonate-bicarbonate buffer pH 9.6. Then, plate is treated with bovine serum albumin solution and washed 4X with PBS-Tween. Then, 50 μl of peroxidase-labeled-protein-AG conjugate diluted 1:3000 in PBS-non-fat milk is added to each well and incubated for 1.30h at RT. After that, the plate is washed 4X with PBS-Tween. Pipette 50 μl of 3,3’,5,5’ - tetramethylbenzidine (TMB; Sigma-Aldrich) to each well. The reaction is stopped with 50 μl of 3M H2SO4 solution. The plate is visually assessed for the development of colour and read in a microplate reader at 450 nm. A cut-off point is calculated as the mean of the optical density of negative controls times two. The cut-off point is 0.28.

Table 34 shows SpAG that binds strongly to 46.66% of the immunoglobulin panel. As compared with SpLG, SpAG interacts with the entire panel except the chicken IgY, and SpLG lacks affinity to IgY of ostriches and chickens and does not interact with the cat IgG.

**Table 34.** SpAG Direct ELISA.

| Immunoglobulins | XOD results | Binding affinity | Coefficient of variation (CV) intra-assay (%) | CV inter-assay (%) |
| --- | --- | --- | --- | --- |
| Mule IgG | 0.42 | + | 3.50 | 5.07 |
| Donkey IgG | 0.44 | + | 4.05 | 7.14 |
| Horse IgG | 0.65 | ++ | 4.15 | 5.90 |
| Dog IgG | 1.12 | +++ | 3.28 | 5.32 |
| Skunk IgG | 1.18 | +++ | 3.58 | 5.78 |
| Coyote IgG | 0.44 | + | 3.67 | 6.60 |
| Raccoon IgG | 1.05 | +++ | 4.06 | 6.81 |
| Ostrich IgY | 0.35 | + | 2.74 | 4.78 |
| Pig IgG | 1.15 | +++ | 4.79 | 7.67 |
| Mouse IgG | 1.09 | +++ | 4.07 | 8.20 |
| Bovine IgG | 0.65 | ++ | 5.13 | 8.05 |
| Rabbit IgG | 1.12 | +++ | 3.76 | 6.03 |
| Chicken IgY | 0.14 | - | 3.43 | 5.79 |
| Cat IgG | 1.19 | +++ | 3.17 | 4.83 |
| Rat IgG | 0.58 | ++ | 3.45 | 5.26 |

### PLAG Direct ELISA

This ELISA is used to study the interactions of protein-LAG (PLAG) with diverse immunoglobulins. The 96 well microtitre plate is coated overnight at 4°C with 1 μg/μl per well of purified immunoglobulins in carbonate-bicarbonate buffer pH 9.6. Then, plate is treated with bovine serum albumin solution and washed 4X with PBS-Tween. Then, 50 μl of peroxidase-labeled-protein-LAG conjugate diluted 1:3000 in PBS-non-fat milk is added to each well and incubated for 1.30h at RT. After that, the plate is washed 4X with PBS-Tween. Pipette 50 μl of 3,3’,5,5’ - tetramethylbenzidine (TMB; Sigma-Aldrich) to each well. The reaction is stopped with 50 μl of 3M H2SO4 solution. The plate is visually assessed for the development of colour and read in a microplate reader at 450 nm. A cut-off point is calculated as the mean of the optical density of negative controls times two. The cut-off point is 0.30.

Table 35 shows a PLAG direct ELISA. This hybrid immunoglobulin-binding protein binds strongly to IgGs from eleven species of animals, which represents 73.33%. In addition, it binds moderately to IgGs of several species as coyote and bovine. This is a very versatile reagent, with the capacity to bind to immunoglobulins from many animal species.

**Table 35.** PLAG Direct ELISA.

| **Immunoglobulins** | **XOD results** | **Binding affinity** | **Coefficient of variation (CV) intra-assay (%)** | **CV inter-assay (%)** |
| --- | --- | --- | --- | --- |
| **Mule IgG** | **0.97** | **+++** | **4.22** | **7.16** |
| **Donkey IgG** | **1.05** | **+++** | **4.34** | **8.38** |
| **Horse IgG** | **0.94** | **+++** | **3.92** | **7.26** |
| **Dog IgG** | **1.00** | **+++** | **4.04** | **7.85** |
| **Skunk IgG** | **1.32** | **+++** | **3.50** | **6.37** |
| **Coyote IgG** | **0.65** | **++** | **4.60** | **7.28** |
| **Raccoon IgG** | **1.25** | **+++** | **3.68** | **6.34** |
| **Ostrich IgY** | **0.49** | **+** | **3.65** | **6.05** |
| **Pig IgG** | **1.45** | **+++** | **4.09** | **7.18** |
| **Mouse IgG** | **1.35** | **+++** | **4.75** | **8.40** |
| **Bovine IgG** | **0.72** | **++** | **4.38** | **6.47** |
| **Rabbit IgG** | **1.18** | **+++** | **4.15** | **7.03** |
| **Chicken IgY** | **0.15** | **-** | **3.65** | **5.70** |
| **Cat IgG** | **1.10** | **+++** | **4.08** | **6.74** |
| **Rat IgG** | **0.95** | **+++** | **3.15** | **4.78** |

The 35 ELISAs were highly reproducible and this is a confirmatory results for some of the ELISAs that have been reported previously as direct (SpL, SpA, SpLA), and sandwich ELISA as SpG-SpLA assay [9,20,25]. The reason why we report that these tests were highly reproducible is the fact that their coefficient of variation (both intra-assays and inter-assays) were within the normal limits except for few immunoglobulin samples. The intra-assay reproducibility was 98.09% (515 tests out of 525 of the samples had CV within the normal limit that were below 5%). The inter-assay reproducibility was 99.62% (meaning that 523 tests out of 525 tests had CV that were below 10%).

Most of the ELISAs are newly reported in this research. They were standardized after a detailed basic ELISA protocol workout. Quantities of proteins being coated in the microplate, washing procedures, optimal sample concentration, optimal dilutions of the conjugates, and optimal molarity of the stop solution reagent were assessed.

It is recommended for further work to test the IgG-binding reagents in ELISAs for immunodetection of zoonotic microorganisms affecting a greater number of mammalian species, as the case of Borrelia burgdorferi. Another suggestion is the demonstration of the binding affinities of immunoglobulin-binding proteins to antibodies of many animal species by Western blot analysis or dot blot analysis.

#### Indirect ELISA for detection of anti-HIV antibodies in cats or rats and dot blot results

Cats, chickens, and rats raised antibodies against a fragment of HIV gp120 that were detected by ELISA and confirmed by dot blot analyses as shown in figures 2 and 3. The authors are not aware of any other documented study describing the use of anti-HIV hyper-immune chicken eggs as oral vaccine. Thibodeau and Tremblay (1991) reported that human immunodeficiency virus (HIV-1) gp160-specific secretory IgA was found in the saliva of rabbits orally immunized with a HIV-immunosome. These antibodies neutralized HIV infectivity in vitro [31].

**Figure 2:**
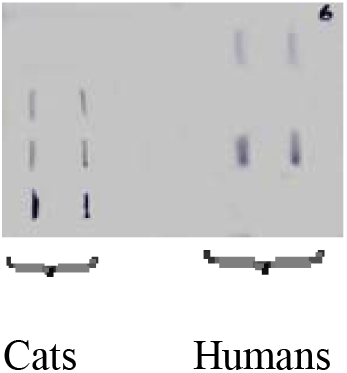
shows dot blotting lines that represent anti-HIV antibodies, which are present in 3 cat sera, orally immunized with anti-HIV hyperimmune chicken eggs. It shows anti-HIV antibodies in the serum from two infected humans (the sera were donated by the National Blood Bank in the Caribbean and it was used as a positive control). No dot blot signals are present in negative controls (sera from two unfed cats and an HIV negative human).

**Figure 3:**
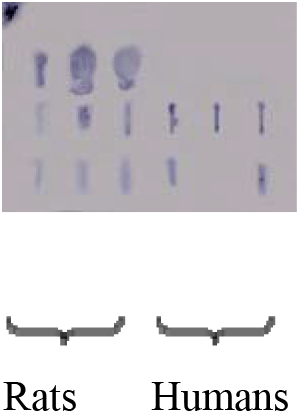
shows 9 dots that represent the presence of anti-HIV antibodies in 3 rats orally immunized with hyperimmune egg raised against the fragment peptide 254-274 of HIV gp120. It shows 5 dots that represent 2 human sera also positive for HIV and were used here as positive controls. The blank spaces represent negative samples or blanks.

Figure 2 shows dot blotting lines that represent anti-HIV antibodies, which are present in 3 cat sera, orally immunized with anti-HIV hyperimmune chicken eggs. It shows anti-HIV antibodies in the serum from two infected humans (the sera were donated by the National Blood Bank in the Caribbean and it was used as a positive control). No dot blot signals are present in negative controls (sera from two un-fed cats and an HIV negative human serum sample).

## Acknowledgements

To the Campus Research and Publication Fund of the University of West Indies, Mona Campus, Jamaica. West Indies. I have gratitude for Professors Norma Anderson and Monica Smikle from University of the West Indies for guidance and support.

## Research Grant

The research was funded by a grant provided by The Caribbean Health Research Council (CHRC) under the CARICOM/EU project on Strengthening the Institutional Response to HIV/AIDS/STIs.

## Conclusions

The single and hybrid immunoglobulin-binding protein were effective in their binding capacity to immunoglobulins from a variety of mammalian species. The potential use of this proteins is in the arena of immunodiagnosis and immunoglobulin detection. Dot blot analysis proves effective in the detection of HIV anti-gp120 antibodies in several animal species including cats, rats, and positive human controls. These antibodies can be used as reagents in the development of immunodiagnostic tests or oral vaccines as the use of hyper-immune chicken eggs.

